# Structural and functional characteristics of SARS-CoV-2 Omicron subvariant BA.2 spike

**DOI:** 10.1101/2022.04.28.489772

**Authors:** Jun Zhang, Weichun Tang, Hailong Gao, Christy L. Lavine, Wei Shi, Hanqin Peng, Haisun Zhu, Krishna Anand, Matina Kosikova, Hyung Joon Kwon, Pei Tong, Avneesh Gautam, Sophia Rits-Volloch, Shaowei Wang, Megan L. Mayer, Duane R. Wesemann, Michael S. Seaman, Jianming Lu, Tianshu Xiao, Hang Xie, Bing Chen

## Abstract

The Omicron subvariant BA.2 has become the dominant circulating strain of severe acute respiratory syndrome coronavirus 2 (SARS-CoV-2) in many countries. We have characterized structural, functional and antigenic properties of the full-length BA.2 spike (S) protein and compared replication of the authentic virus in cell culture and animal model with previously prevalent variants. BA.2 S can fuse membranes more efficiently than Omicron BA.1, mainly due to lack of a BA.1-specific mutation that may retard the receptor engagement, but still less efficiently than other variants. Both BA.1 and BA.2 viruses replicated substantially faster in animal lungs than the early G614 (B.1) strain in the absence of pre-existing immunity, possibly explaining the increased transmissibility despite their functionally compromised spikes. As in BA.1, mutations in the BA.2 S remodel its antigenic surfaces leading to strong resistance to neutralizing antibodies. These results suggest that both immune evasion and replicative advantage may contribute to the heightened transmissibility for the Omicron subvariants.

## Introduction

The next wave of COVID-19 (coronavirus disease 2019) cases will probably be driven by new Omicron subvariants of severe acute respiratory syndrome coronavirus 2 (SARS-CoV-2), as the Omicron BA.2 viruses have gradually replaced the dominant BA.1 strain in many countries (*1, 2*). These viruses are generally believed to be milder than the previously dominant variants of concern (VOCs), but still have caused a high death rate in Hong Kong, particularly among unvaccinated elderly (*3, 4*). BA.2 infection appears to produce somewhat higher viral loads in humans than that by BA.1 (*1*), consistent with the data from cell-culture experiments showing that BA.2 replicates more efficiently in nasal epithelial cells and is more fusogenic than BA.1 (*5*). Nevertheless, the clinical outcomes of infection by these two Omicron subvariants do not appear to be very different (*6*). Both BA.1 and BA.2 are antigenically distant from the Wuhan-Hu-1 stain that initiated the pandemic (*7, 8*), and they evade neutralizing antibodies elicited by the Wuhan-Hu-1 based vaccines or by infection of early viral strains to a similar degree (*9*), but with some qualitative differences (*10, 11*). Vaccinated individuals infected with BA.1 can induce cross-reactive immunity to BA.2, and reinfection with BA.2 after BA.1 infection is rare (*12–14*). Protective efficacy by the first-generation vaccines against symptomatic COVID-19 after booster is also similar for BA.1 and BA.2 (*15, 16*). It remains important to understand the molecular characteristics of the BA.2 subvariant for developing effective intervention strategies.

The SARS-CoV-2 spike (S) protein is the main component of all COVID-19 vaccines (*17*). Its primary function is to catalyze fusion of the viral and target cell membranes, upon engaging the cellular receptor angiotensin converting enzyme 2 (ACE2), to allow the viral RNA genome to enter a host cell and initiate infection. The protein is produced as a singlechain precursor and cleaved by a host furin-like protease into the receptor-binding fragment S1 and the fusion fragment S2 (Fig. S1; ref(*18*)). The mature viral spike is formed by three copies of noncovalently-associated S1 and S2 fragments, and induces strong host immune responses during natural infection. After binding of S1 to the receptor, ACE2, on the host cell, S2 is further cleaved at the S2’ site (Fig. S1) by either TMPRSS2 (transmembrane serine protease 2) or cathepsins L (*19*), inducing dissociation of S1 and a series of conformational changes in S2, and ultimately the fusion of viral and cell membranes (*20–22*). S1 contains four domains – the N-terminal domain (NTD), receptor-binding domain (RBD), and two CTDs (C-terminal domains). Three copies of S1 wrap around the central helical-bundle structure formed by the prefusion S2 trimer. The RBD can adopt either a “down” conformation for a receptor-inaccessible state that protects its receptor binding site from the host immune system, or an “up” conformation for a receptor-accessible state for effective ACE2 engagement (*23, 24*). Like BA.1, which acquired an unusually large number of mutations in its S protein, the BA.2 spike also has ~30 residue changes, as compared to 10 mutations on average in other VOCs (*25*). The main differences in S between BA.1 and BA.2 are in its NTD, with no shared common mutations. BA.2 has three “new” mutations (T376A, D405N and R408S) with respect to BA.1 in the RBD, but lacks two (G446S and G496S) in the RBD and an additional two (N856K and L981F) in the S2 region.

In this study, we have characterized the full-length S protein of the Omicron BA.2 subvariant and determined its structures with a native sequence by cryogenic electron microscopy (cryo-EM). Comparison of the structure, function and antigenicity of the BA.2 S, as well as replication of the authentic viruses in cell culture and animal model, with those of the previously characterized VOCs has provided molecular insights into this highly transmissible SARS-CoV-2 variant.

## Results

### BA.2 S fuses membrane more efficiently than BA.1 but less than other VOCs

To assess the fusogenic activity of the full-length BA.2 S protein derived from a natural isolate (hCoV-19/Denmark/DCGC-327158/2022; Fig. S1), we first expressed the protein in HEK293 cells and compared its membrane fusion activity with that of the full-length S constructs from previous variants, including G614, Alpha, Beta, Delta and Omicron-BA.1 (*26–29*). All S proteins expressed at comparable levels, with similar degrees of furin cleavage for G614, BA.1 and BA.2, but slightly more for Beta, Alpha and Delta at 24 hours posttransfection (Fig. S2A). Under our standard cell-cell fusion assay conditions (*30*), the cells producing these S proteins fused efficiently with ACE2-expressing cells, except that the fusion activities of both the BA.1 and BA.2 S proteins were somewhat lower than those of other S proteins, including G614 (Fig. S2B).

We next performed a time-course experiment with both S and ACE2 expressed at saturating levels (Fig. 1A; ref (*30*)). Both Omicron BA.1 and BA.2 S proteins again had consistently lower activities than other variants, but they were almost indistinguishable from each other throughout the time period tested (Fig. 1A). When untransfected HEK293 cells expressing a minimal level of endogenous ACE2 were used, all other variants had significant fusion activities, particularly at later time points, while BA.1 S was essentially inactive, as reported previously (*29*), and BA.2 had a barely detectable activity only at the last time point (Fig. S3A), suggesting that the two Omicron subvariants may not infect host cells with very low levels of ACE2. We then analyzed how these S proteins responded to different amounts of ACE2 expressed in HEK293 cells. The fusion activities increased with the increased level of ACE2 for all variants, as expected, but both BA.1 and BA.2 lagged the other variants and required higher levels of ACE2 to reach a similar fusion activity, until they plateaued at the saturating ACE2 level (Fig. 1B). We note that BA.2 was significantly more fusogenic than BA.1 at mid-range ACE2 levels. Likewise, the replication of both the authentic BA.1 and BA.2 viruses in VeroE6 cells (*31*) was also slower than that of the G614 and Delta viruses at least within the first 48 hours postinfection (Figs. 1C and S4). In another time-course experiment using murine leukemia virus (MLV)-based pseudoviruses expressing the S constructs with the cytoplasmic tail deleted (*28*), Delta and Delta AY.4 subvariant infected target cells expressing a high level of ACE2 much more rapidly than other variants tested, as reported previously (*28, 29*), while both BA.1 and BA.2 pseudoviruses showed kinetics of infection similar to those of G614 and Alpha variants (Fig. S3B). These results suggest that the BA.2 S has a slight advantage in its membrane fusion capability than BA.1, but remains a slightly weakened fusogen when compared to other VOCs.

**Figure 1.**
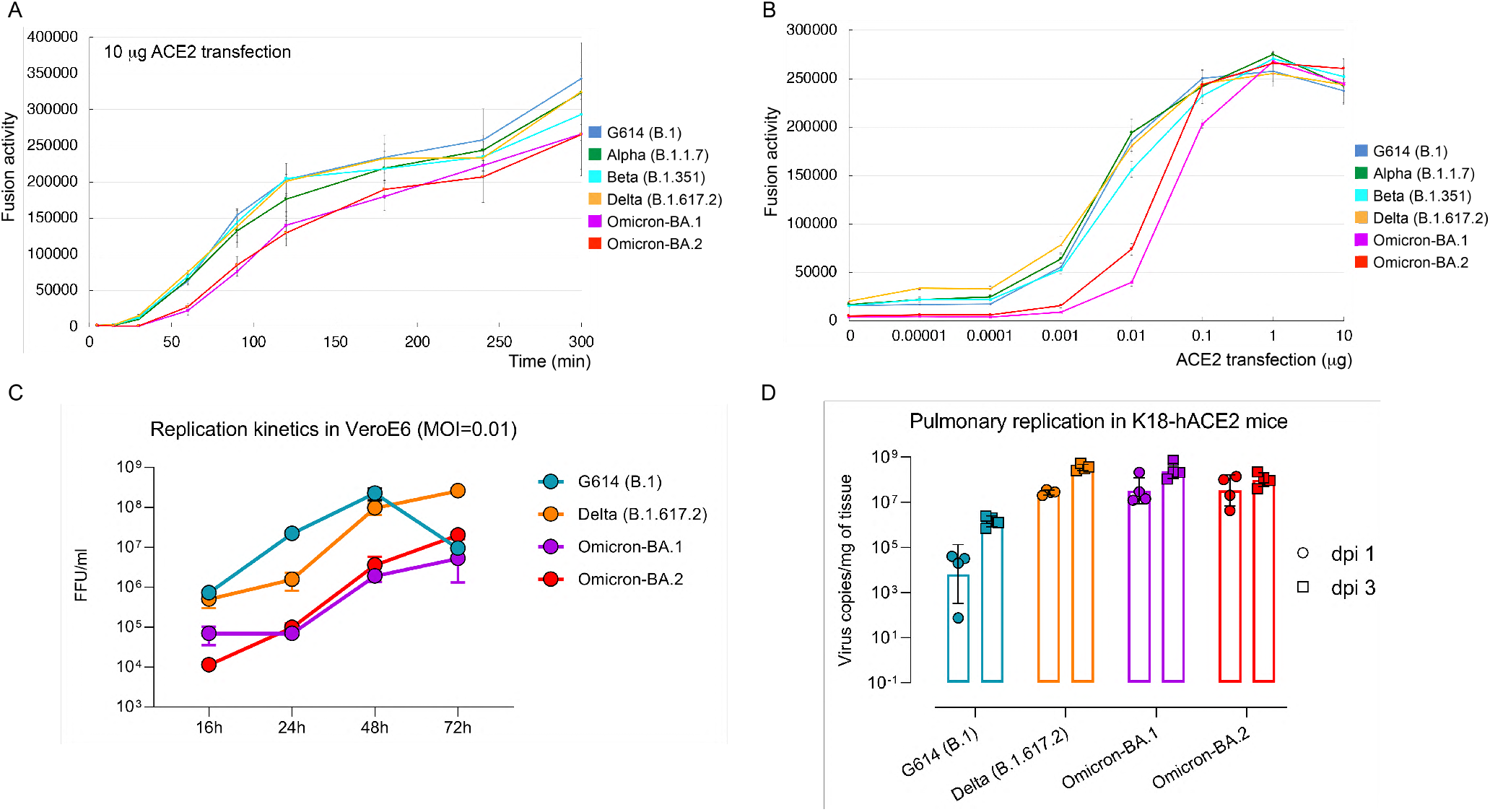
Functional properties of the Omicron BA.2 spike. (**A**) Time-course of cellcell fusion mediated by various full-length S proteins, as indicated, with the target HEK293 cells transfected with 10 μg ACE2. (**B**) Cell-cell fusion mediated by various full-length S proteins with HEK293 cells transfected with various levels (0-10 μg) of the ACE2 expression construct. (**C**) Authentic virus replication kinetics in Vero E6 cells. Focus-forming units (FFU) per ml were determined and data are expressed as mean±sem (n=3 independent replicates/time point/virus). The foci images of in vitro replication kinetics of these viruses are shown in Fig. S4. (**D**) Pulmonary replication in K18-hACE2 mice at 1 and 3 days post infection (dpi). Mice were intranasally inoculated with 100 TCID_50_/mouse of authentic G614 (B.1), Delta (B.1617.2), Omicron-BA.1 or BA.2 viruses. Viral RNA copies in whole lung homogenates were measured by RT-qPCR (n=4 mice/time point/group). Data are expressed as geometric means (bars) with geometric standard deviation (error bars) and individual pulmonary viral burdens are shown.

### BA.1 and BA.2 are fast replicating viruses in K18-hACE2 transgenic mice

We compared the replication of the authentic G614, Delta, Omicron BA.1 and BA.2 viruses in K18-hACE2 transgenic mice, which express human ACE2 and are susceptible to SARS-CoV-2 (*32, 33*). All K18-hACE2 mice were genotype-confirmed, randomly grouped, and inoculated intranasally with each of the selected viruses at 100 TCID_50_/mouse under isoflurane anesthesia. Viral loads were determined in various tissues, including brain, heart, nasal turbinates, whole lung, liver, spleen and kidneys, harvested from euthanized animals at 1 or 3 days post infection (dpi). As shown in Figs. 1D and S5, these tissues expressed human ACE2 at different levels, with the nasal turbinates among the highest, the heart the lowest and the lung in-between. The human ACE2 expression in these mouse tissues was not altered significantly during the early stage of viral infection. Intranasal delivery under anesthesia generally forced the viruses to pass through the nasal passage and to reach the lung of the inoculated animals directly without first colonization of the upper respiratory tract. Not surprisingly, the lung tissues, where the primary infection began, showed the highest viral loads in all infected mice. Viral dissemination to extrapulmonary organs showed no apparent correlation with the human ACE2 expression levels in these mice. The viral loads in lung tissues with Delta, BA.1 or BA.2 infection were 2-3 orders of magnitude higher than those infected by the G614 virus, suggesting that both BA.1 and BA.2 are fast replicating viruses, similar to Delta, at least in this animal model, despite the compromised fusogenicity of their spike proteins. Therefore, the heightened transmissibility of the Omicron variants does not appear to require increased viral entry.

### Biochemical and antigenic properties of the intact BA.2 S protein

To produce full-length BA.2 S protein with no stabilizing modifications, we transfected HEK293 cells with a C-terminal strep-tagged construct (Fig. S6A), and purified the protein following a protocol established for other intact S protein preparations (*26, 27, 29, 30*). Resolved by gel-filtration chromatography, the purified Wuhan-Hu-1 S protein elutes in three peaks, corresponding to the prefusion S trimer, postfusion S2 trimer and dissociated S1 monomer, respectively; the G614 S protein shows a single peak for the prefusion trimer (Fig. S6B; ref (*26, 30*)). The gel-filtration profile of BA.2 protein was almost identical to that of the BA.1 protein (*29*), eluting in one major peak corresponding to the prefusion trimer but with a significant amount of aggregate on the leading side and also a shoulder on the trailing side (Fig. S6B). SDS-PAGE analysis showed that the BA.2 protein was cleaved significantly less than was the G614 trimer at 84 hours posttransfection (Fig. S6B), indicating that furin cleavage of the BA.2 subvariant, like that of BA.1 (*29*), is inefficient, and that the two mutations (N679K and P681H) in the Omicron S proteins near the furin site have reduced its susceptibility to the proteolytic cleavage.

We analyzed receptor-binding and antigenic properties of the prefusion BA.2 S trimer by comparing its binding to soluble ACE2 proteins and anti-S monoclonal antibodies with that of the G614 trimer using bio-layer interferometry (BLI). Competition assays with an unbiased panel of antibodies isolated from COVID-19 convalescent individuals have defined seven epitopic regions on the S trimer, designated RBD-1, RBD-2, RBD-3, NTD-1, NTD-2, S2-1 and S2-2 (Fig. S7A; ref(*34*)). The BA.2 variant bound substantially more strongly to both the ACE2 monomer or dimer than did the G614 trimer, but only slightly more than BA.1 (Figs. 2 and S7B; Table S1; ref (*29*)), suggesting that the differences in the mutations of the RBD between BA.1 and BA.2 have little impact on these relative receptor binding affinities.

**Figure 2.**
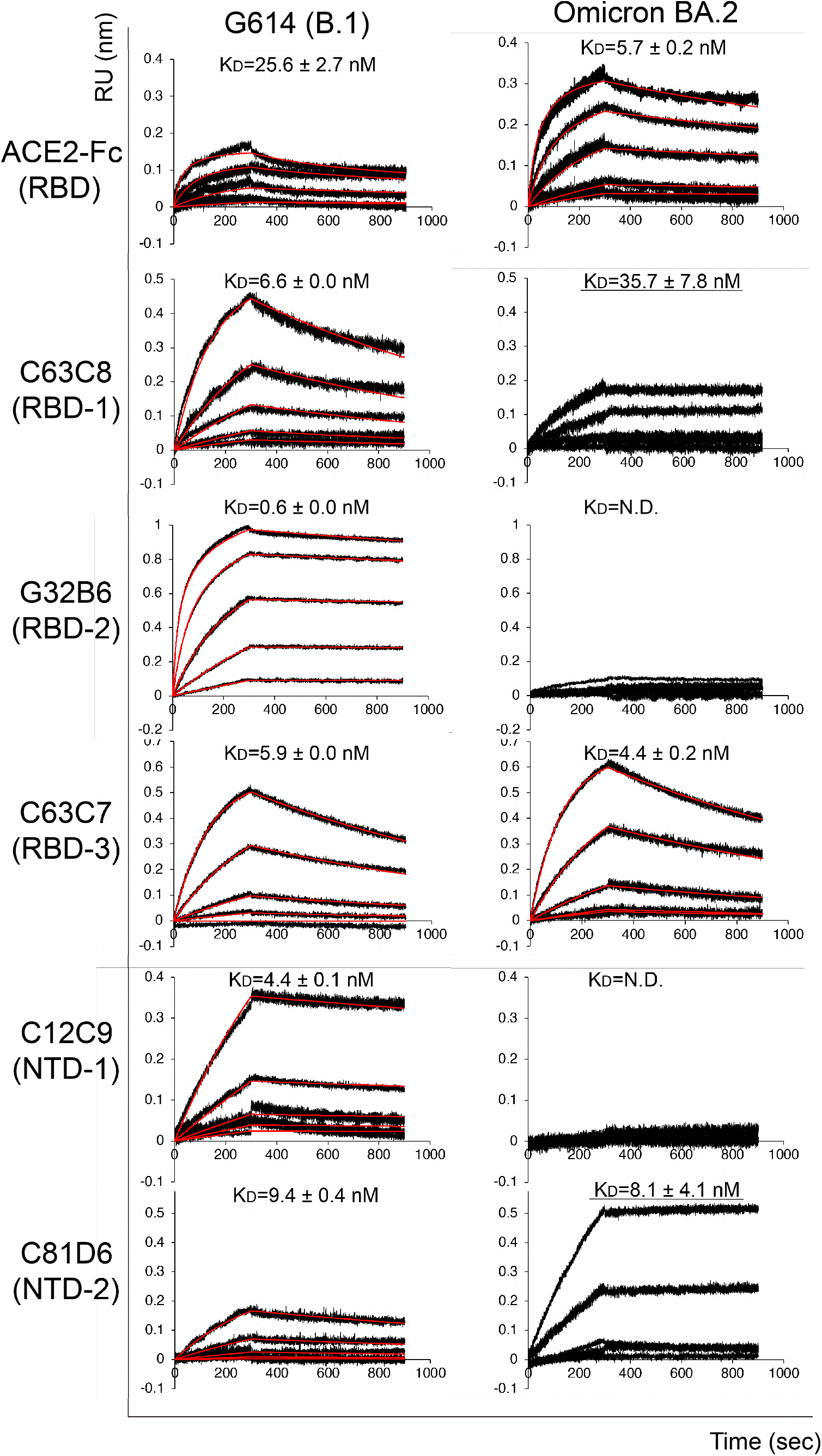
Antigenic properties of the purified full-length BA.2 S protein. Bio-layer interferometry (BLI) analysis of the association of prefusion S trimers derived from the parental G614 strain and Omicron BA.2 subvariant with a soluble dimeric ACE2 construct and with a panel of antibodies targeting five epitopic regions on the RBD and NTD (see Fig. S6A and ref(*34*)). For ACE2 binding, the purified ACE2 protein was immobilized to AR2G biosensors and dipped into the wells containing each purified S protein at various concentrations. For antibody binding, various antibodies were immobilized to AHC biosensors and dipped into the wells containing each purified S protein at different concentrations. Binding kinetics were evaluated using a 1:1 Langmuir model except for dimeric ACE2 and antibody G32B6 targeting the RBD-2, which were analyzed by a bivalent binding model. The sensorgrams are in black and the fits in red. Binding constants highlighted by underlines were estimated by steady-state analysis as described in the Methods. RU, response unit. Binding constants are also summarized here and in Table S1. N.D., not determined. All experiments were repeated at least twice with essentially identical results.

We reported previously that selected monoclonal antibodies from the unbiased panel bind strongly to the G614 S trimer (*27–29*). BA.2 lost binding completely to the two NTD-1 antibodies, 12C9 and C83B6, and the two RBD-2 antibodies, G32B6 and C12A2, and bound less strongly than G614 S to the RBD-1 antibodies, C63C8 and G32R7, as well as to the nonneutralizing S2 antibody 163E6 (Figs. 2 and S7B; Table S1). Its affinity for the non-neutralizing NTD-2 antibody C81D6 appeared to be higher than that of the G614 trimer. These BLI data were largely consistent with binding of these antibodies to membrane-bound S trimers measured by flow cytometry (Fig. S8). We determined, in an HIV-based pseudovirus assay, the neutralization potencies of these antibodies and of additional selected ones for each group (*34*), as well as that of a designed trimeric ACE2 (*35*), against infection by the BA.2 subvariant. As summarized in Table S2, BA.2, like BA.1, was almost completely resistant to all NTD-1, NTD-2 and RBD-3 antibodies, as well as RBD-2 antibodies except for C98C7, which had much reduced potency. BA.2 appeared to retain some weak sensitivities to RBD-1 antibodies, slightly different from the pattern observed for BA.1 (Table S2). The trimeric ACE2 is significantly more potent against the two Omicron subvariants than G614, consistent with their greater receptor binding affinity. Overall, the neutralization data are largely in agreement with their binding affinity for the membrane-bound or purified S proteins.

### Cryo-EM structure of the full-length BA.2 S trimer

We determined the cryo-EM structures of the full-length BA.2 S trimer expressed using an unmodified sequence derived from a natural isolate. We recorded cryo-EM images on a Titan Krios electron microscope equipped with a Gatan K3 direct electron detector, and used cryoSPARC (*36*) for particle picking, two-dimensional (2D) classification, threedimensional (3D) classification and refinement (Fig. S9). Three distinct classes were obtained from 3D classification, representing the closed, three-RBD-down prefusion conformation, a one-RBD-up conformation, and an RBD-intermediate conformation, all three also found for the G614 trimer (*26*). These classes were further refined to 3.0-3.5Å resolution (Fig. S9-S12; Table S3). There are no major differences in overall architecture between the BA.2 S protein and the G614 S trimer in the corresponding conformation (Fig. 3A and 3B; ref(*26*)). Additional structured residues near the furin cleavage site at the S1/S2 boundary, present in the BA.1 structures (*29*), were also present in BA.2. We previously suggested that the sidechain of Lys679, resulting from the N679K mutation, could make contacts with the nearby CTD-2 and rigidify the furin cleavage loop, which is largely disordered in the previous variants (Fig. S13). Thus, the more structured cleavage site may retard its docking into the furin active site, possibly explaining why the cleavage efficiency of the two Omicron S proteins is lower than that of other variants.

**Figure 3.**
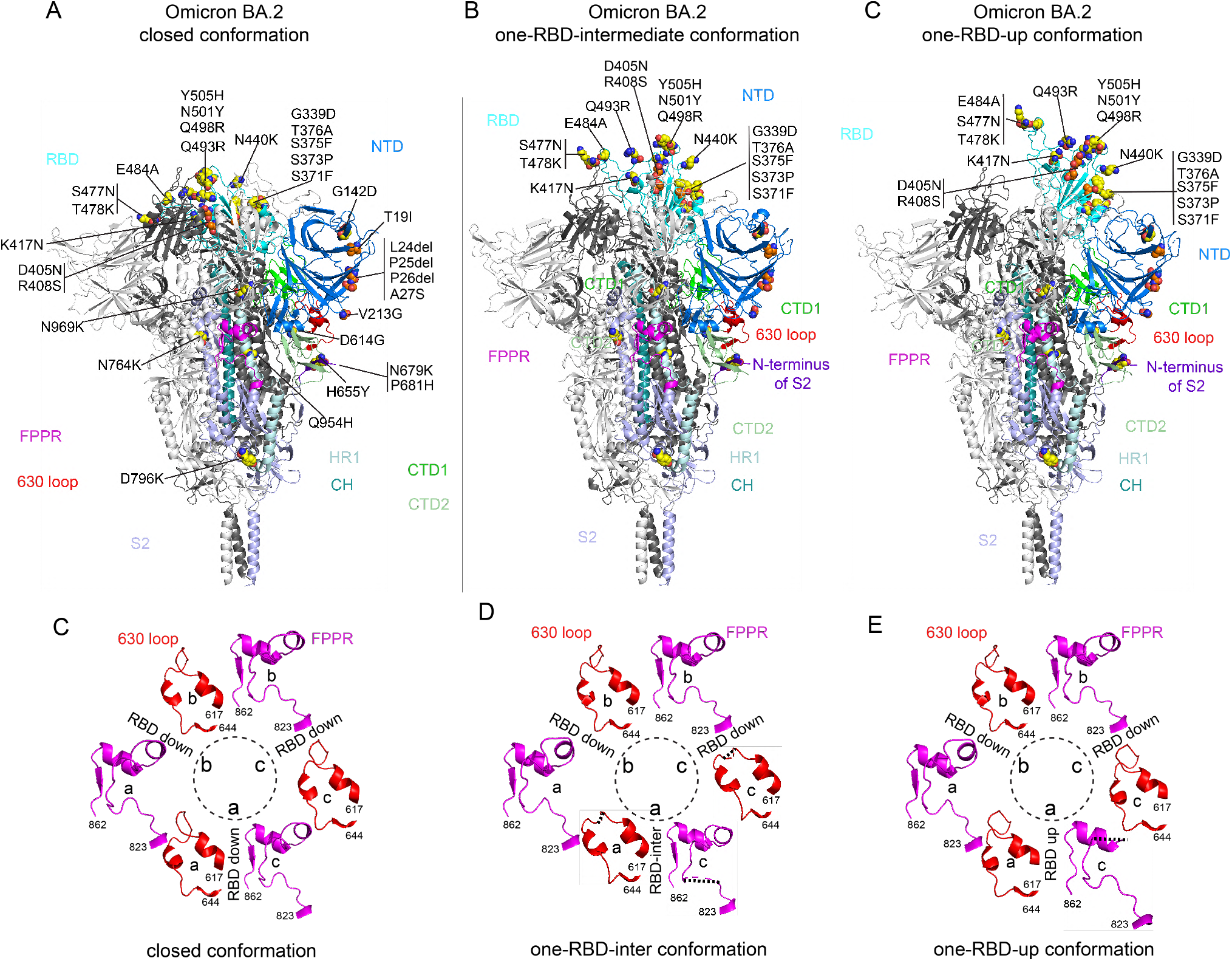
Cryo-EM structures of the full-length Omicron S protein. (**A)-(C**) The structures of the closed prefusion conformation, one-RBD-intermediate and one-RBD-up conformations of the full-length BA.2 S trimer, respectively, are shown in ribbon diagram with one protomer colored as NTD in blue, RBD in cyan, CTD1 in green, CTD2 in light green, S2 in light blue, the 630 loop in red, FPPR in magenta, HR1 in light blue, CH in teal and the N-terminal segment of S2 in purple. All BA.2 mutations, as compared to the Wuhan-Hu-1 sequence, are highlighted in sphere model, with those unique to BA.2 colored in orange. (**C**)-(**E**) Structures, in the BA.2 closed, one-RBD-intermediate and one-RBD-up conformations, respectively, of segments (residues 617-644) containing the 630 loop (red) and segments (residues 823-862) containing the FPPR (magenta) from each of the three protomers (a, b and c). The position of each RBD is indicated. Dashed lines indicate gaps in the chain trace (disordered regions).

Our studies on the previous VOCs have identified the FPPR (fusion peptide proximal region) and 630 loop as two control elements that may modulate the RBD movement and the S protein stability (*26, 30, 37*). They have to move away from their original positions in the closed conformation when the RBD flip up to expose the receptor binding site, and they become at least partially disordered in the cryo-EM maps for the RBD-up conformations. In the BA.1 S trimer, the mutation N856K has led to a new salt bridge between the FPPR (Lys856) and the CTD-1 (Asp568) of the neighboring protomer, and stabilized the FPPR even when the RBD above it move to an up-conformation. Indeed, the flipped RBD moves further up to avoid clashes with the FPPR underneath, creating additional kinetic barriers for the RBD down-to-up transition and thus for ACE2 engagement (*29*). In the three-RBD-down conformation of the BA.2 trimer (Fig. 3C), the FPPRs and 630 loops are all structured, as expected. In the one-RBD-intermediate or one-RBD-up conformation of the G614 trimer, only one FPPR and 630-loop pair is ordered (*26*). Two FPPRs and one 630 loop in the one-RBD-intermediate BA.2 trimer were structured, while one FPPR became partially disordered in the one-RBD-up conformation (Fig. 3C-D). The mutation N856K is not present in BA.2, but apparently it is not the only structural constraint on these elements, and its absence does not free the RBD movement seen in the other variants.

### Structural impact of the BA.2 mutations

The overall structure of the BA.2 RBD was almost identical to that of BA.1, with the same small shift of a short helix (residues 365-371; Fig. 4A) found in BA.1 (*29*). The ACE2 binding interface and many antibody epitopes from the RBD-2 group (Fig. S7A) were similarly altered by the large number of surface mutations in the BA.2 RBD. N501Y, Q493R and Q498R have been shown to enhance ACE2 affinity by creating additional contacts with the receptor (*38–42*), while K417N and E484A eliminate ionic interactions with ACE2 and may reduce the ACE2 affinity (*27, 43*). All these mutations are found in both BA.1 and BA.2, and may be responsible for the observed increase in receptor affinity (*39, 40, 42*). The unique mutations in the BA.2 RBD – D405N, R408S and T376A – are located outside the ACE2 binding site, and they therefore had minimal impact on receptor binding. Most mutations, some shared by BA.1 and BA.2, are at the periphery of the epitopes targeted by the RBD-1 and RBD-3 antibodies (Fig. S14), explaining why their impact on binding and neutralization of these antibodies is largely antibody-dependent (Table S1 and S2). In contract, mutations K417N, S477N, T478K, E484A, Q493R, Q498R and N501Y are present in both BA.1 and BA.2 at the ACE2 binding interface, and these residues form part of the major contacts with RBD-2 antibodies, accounting for complete loss of binding and neutralization of the two Omicron subvariants by most RBD-2 antibodies. Thus, the RBD-2 region, heavily involved in receptor binding, has greater tolerance to surface mutations than the RBD-1 and RBD-3 regions.

**Figure 4.**
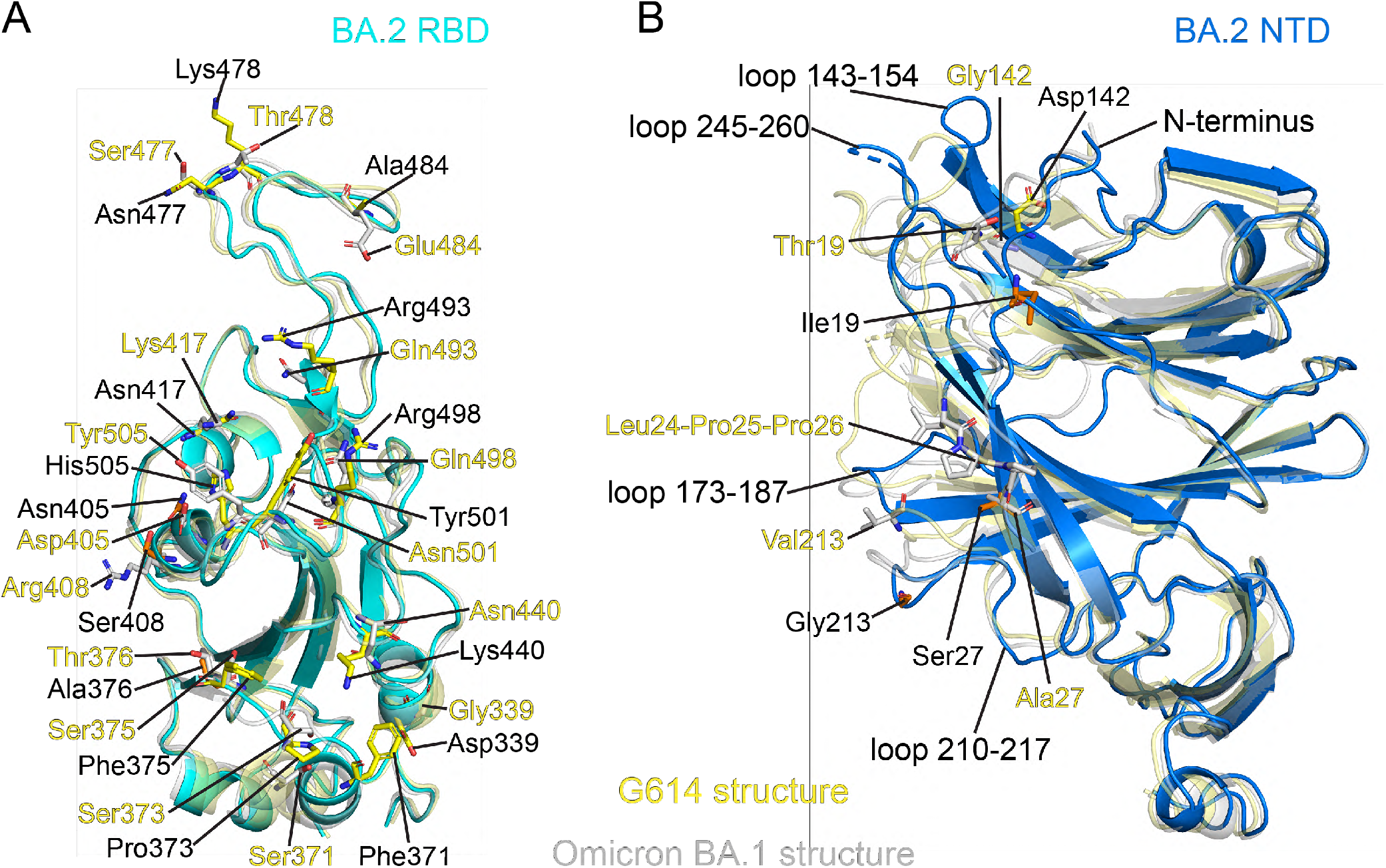
Structural impact of the mutations in the BA.2 S. (**A**) Superposition of the RBD structure of the BA.2 S trimer in cyan with the RBD of the G614 S trimer in yellow and the RBD of the BA.1 S trimer in gray. Locations of all 16 BA.2 mutations in the RBD are indicated and these residues are shown in stick model with those unique to BA.2 in orange. (**B**) Superposition of the NTD structure of the BA.2 S trimer in blue with the NTD of the G614 S trimer in yellow and the NTD of the BA.1 S trimer in gray. Locations of mutations T19I, A27S, G142D, V213G and the deletion L24del-P25del-P26del are indicated and these residues are shown in stick model with those unique to BA.2 in orange. The N-terminal segment, 143-154, 173-187, 210-217 and 245-260 loops are rearranged between the two structures.

The mutation S373P, which may rigidify the local polypeptide chain, present in both BA.1 and BA.2, causes a shift of the short helix formed by residues 365-371, first observed in the BA.1 RBD (*29*). The shift apparently forced a large rotation of the N-linked glycan at Asn343, with the side chain of the mutated residue 371 (S371L in BA.1 and S371F in BA.2) projecting towards the reconfigured glycan (Figs. 4B and S14).

The BA.2 NTD has four point mutations (T19I, A27S, G142D and V213G) and one three-residue deletion (L24del-P25del-P26del), all of which are different from those found in BA.1 and apparently sufficient to reconfigure the N-terminal segment and the surface-exposed 143-154, 173-187, 210-217 and 245-260 loops, several of which form important parts of the neutralizing epitopes in the NTD (*44*). Thus, these structural changes, substantially different from those found in the previous variants, have nevertheless altered the antigenic surface of the NTD enough to account for the loss of binding and neutralization by NTD-1 antibodies (Figs. 2 and S6; Table S1 and S2).

The remaining mutations in BA.2, including N764K, D796Y, Q954H and N969K, all shared with BA.1, did not lead to any obvious structural changes, as also observed in the BA.1 structures (*29*).

## Discussion

SARS-CoV-2 variants of concern are generally selected for their transmissibility and the ability to evade host immunity. For the previous COVID-19 waves, the Alpha and Delta variants outcompeted the circulating viruses at the time primarily by their enhanced transmissibility when a large fraction of the population had yet to gain immunity from either natural infection or vaccination, and when acquired immunity had begun to wane (*28, 45–48*). Subsequently, the Omicron lineages emerged with a much larger number of mutations in the spike gene than the previous VOCs and its BA.1 subvariant rapidly supplanted Delta, followed by a gradual replacement of the apparently more contagious BA.2 subvariant (*1, 2*). In both the BA.1 and BA.2 subvariants, the mutations in the spikes not only have caused large rearrangements of the antigenic structure of the NTD, but have also altered the antigenic surface of the RBD considerably, leading to the strong resistance to neutralizing antibodies at a level not seen in any other early variants. Paradoxically, many studies find that the excessive number of mutations in BA.1 S protein may have compromised its fusogenic capability in exchange for its ability to evade host immunity despite its greater ACE2 binding affinity (*49–55*). We further showed that BA.1 S required a substantially higher level of ACE2 on the host cells for efficient membrane fusion, mainly because the mutation N856K stabilizes the FPPR and may retard an order-disorder transition in the FPPR, thereby introducing additional kinetic barriers for the RBD “down” to “up” transition that exposes the receptor binding site (*29*). This mutation is not found in the BA.2 S, which is indeed more fusogenic within a range of ACE2 levels than BA.1, although its spike is still a weaker fusogen than those of other VOCs. Thus, the data presented here provided a molecular basis for the greater transmissibility of BA.2 than that of BA.1 (*2*).

Our study with the K18-hACE2 mice infected by the authentic viruses have also given some clues for why Omicron subvariants are more contagious than the G614 (B.1) strain. These transgenic mice express the human ACE2 under a cytokeratin 18 promoter (K18) and are highly susceptible to SARS-CoV-2 infection, which can result in tissue pathology, inflammatory responses, and dose-dependent disease courses, mimicking the COVID-19 in humans (*33, 56*). As the intranasal delivery forces the viruses directly into lung, we found that pulmonary viral RNA copy numbers there were 100-1000 fold higher within the first 24 hours postinfection in animals infected with Delta and Omicron variants than in those infected with the wildtype G614 virus. Thus, before host immune responses are mounted, both BA.1 and BA.2 can replicate in lung of susceptible animals as rapidly as does the Delta variant and substantially more rapidly than the G614 virus, despite their compromised viral entry, demonstrated in cell culture. We note that the ability of these viruses to disseminate into extrapulmonary organs does not correlate with the receptor ACE2 levels and is likely influenced by other host factors and/or postinfection responses. Together these results suggest that both immune evasion by the spike protein and replicative advantage by the replication machinery (*57*) may contribute to the heightened transmissibility for the Omicron subvariants.

The BA.2 subvariant has evolved a strategy distinct from those seen previously to remodel the NTD with just 7 residue changes. It preserves the overall structure of the RBD, despite 16 point mutations, because of the functional importance of the receptor binding. It is intriguing that almost all the RBD mutations are located along the edges of the RBD-1 and RBD-3 epitopic regions, while there are multiple changes in the center of the RBD-2 region that directly overlaps with the ACE2 binding site. Because RBD-1 and RBD-3 surfaces have been largely conserved, targeting the center of the RBD-1 or RBD-3 regions, clearly immunogenic, may be a promising strategy for immunogen design for inducing broadly neutralizing antibody responses that can protect against current and perhaps future variants of SARS-CoV-2.

## Materials and Methods

### Expression constructs

The gene of a full-length spike (S) protein from Omicron BA.2 variant (hCoV-19/Denmark/DCGC-327158/2022; GISAID accession ID: EPI_ISL_9070870) was assembled from DNA fragments synthesized by Integrated DNA Technologies, Inc. (Coralville, Iowa). The S gene was fused with a C-terminal twin Strep tag (SGGGSAWSHPQFEKGGGSGGGSGGSSAWSHPQFEK) and cloned into a mammalian cell expression vector pCMV-IRES-puro (Codex BioSolutions, Inc, Gaithersburg, MD).

### Expression and purification of recombinant proteins

Expression and purification of the full-length S proteins were carried out as previously described (*30*). Briefly, expi293F cells (ThermoFisher Scientific, Waltham, MA) were transiently transfected with the S protein expression constructs. To purify the S protein, the transfected cells were lysed in a solution containing Buffer A (100 mM Tris-HCl, pH 8.0, 150 mM NaCl, 1 mM EDTA) and 1% (w/v) n-dodecyl-β-D-maltopyranoside (DDM) (Anatrace, Inc. Maumee, OH), EDTA-free complete protease inhibitor cocktail (Roche, Basel, Switzerland), and incubated at 4°C for one hour. After a clarifying spin, the supernatant was loaded on a strep-tactin column equilibrated with the lysis buffer. The column was then washed with 50 column volumes of Buffer A and 0.3% DDM, followed by additional washes with 50 column volumes of Buffer A and 0.1% DDM, and with 50 column volumes of Buffer A and 0.02% DDM. The S protein was eluted by Buffer A containing 0.02% DDM and 5 mM d-Desthiobiotin. The protein was further purified by gel filtration chromatography on a Superose 6 10/300 column (GE Healthcare, Chicago, IL) in a buffer containing 25 mM Tris-HCl, pH 7.5, 150 mM NaCl, 0.02% DDM. All RBD proteins were purchased from Sino Biological US Inc (Wayne, PA).

The monomeric ACE2 or dimeric ACE2 proteins were produced as described (*35*). Briefly, Expi293F cells transfected with monomeric ACE2 or dimeric ACE2 expression construct and the supernatant of the cell culture was collected. The monomeric ACE2 protein was purified by affinity chromatography using Ni Sepharose excel (Cytiva Life Sciences, Marlborough, MA), followed by gel filtration chromatography. The dimeric ACE2 protein was purified by GammaBind Plus Sepharose beads (GE Healthcare), followed gel filtration chromatography on a Superdex 200 Increase 10/300 GL column. All the monoclonal antibodies were produced as described (*34*).

### Western blot

Western blot was performed using an anti-SARS-COV-2 S antibody following a protocol described previously (*58*). Briefly, full-length S protein samples were prepared from cell pellets and resolved in 4-15% Mini-Protean TGX gel (Bio-Rad, Hercules, CA) and transferred onto PVDF membranes. Membranes were blocked with 5% skimmed milk in PBS for 1 hour and incubated a SARS-CoV-2 (2019-nCoV) Spike RBD Antibody (Sino Biological Inc., Beijing, China, Cat: 40592-T62) for another hour at room temperature. Alkaline phosphatase conjugated anti-Rabbit IgG (1:5000) (Sigma-Aldrich, St. Louis, MO) was used as a secondary antibody. Proteins were visualized using one-step NBT/BCIP substrates (Promega, Madison, WI).

### Cell-cell fusion assay

The cell-cell fusion assay, based on the α-complementation of E. coli β-galactosidase, was used to measure fusion activity of SARS-CoV2 S proteins, as described (*30*). Briefly, to produce S-expressing cells, HEK293T cells were transfected by polyethylenimine (PEI; 80 μg) with either 5 or 10 μg of the full-length SARS-CoV2 (G614, Alpha, Beta, Delta or Omicron) S construct, as indicated in each specific experiment, and the α fragment of E. coli β-galactosidase construct (10 μg), as well as the empty vector to make up the total DNA amount to 20 μg. To create target cells, the full-length ACE2 construct at various amount (10 pg-10 μg), as indicated in each specific experiment, the ω fragment of E. coli β-galactosidase construct (10 μg) and the empty vector were used to transfect HEK293T cells. After incubation at 37°C for 24 hrs, the cells were detached using PBS buffer and resuspended in complete DMEM medium. 50 μl S-expressing cells (1.0×10^6^ cells/ml) were mixed with 50 μl ACE2-expressing target cells (1.0×10^6^ cells/ml) to allow cell-cell fusion to proceed at 37°C for 2 hrs for our standard assays or from 10 min to 5 hours for the timecourse experiments. Cell-cell fusion activity was quantified using a chemiluminescent assay system, Gal-Screen (Applied Biosystems, Foster City, CA), following the standard protocol recommended by the manufacturer. The substrate was added to the cell mixture and allowed to react for 90 min in dark at room temperature. The luminescence signal was recorded with a Synergy Neo plate reader (Biotek, Winooski, VT).

### Binding assay by bio-layer interferometry (BLI)

Binding of monomeric or dimeric ACE2 to the full-length Spike protein of each variant was measured using an Octet RED384 system (ForteBio, Fremont, CA). Briefly, monomeric ACE2 or dimeric ACE2 was immobilized to Amine Reactive 2nd Generation (AR2G) biosensors (ForteBio, Fremont, CA) and dipped in the wells containing the S protein at various concentrations (G614 or BA.2, 0.617-150 nM) for association for 5 minutes, followed by a 10 min dissociation phase in a running buffer (PBS, 0.02% Tween 20, 0.02% DDM, 2 mg/ml BSA). To measure binding of a full-length S protein to monoclonal antibodies, the antibody was immobilized to anti-human IgG Fc Capture (AHC) biosensor (ForteBio, Fremont, CA) following a protocol recommended by the manufacturer. The full-length S protein was diluted using a running buffer (PBS, 0.02% Tween 20, 0.02% DDM, 2 mg/ml BSA) to various concentrations (0.617-50 nM) and transferred to a 96-well plate. The sensors were dipped in the wells containing the S protein solutions for 5 min, followed with a 10 min dissociation phase in the running buffer. Control sensors with no ACE2 or antibody were also dipped in the S protein solutions and the running buffer as references. Recorded sensorgrams with background subtracted from the references were analyzed using the software Octet Data Analysis HT Version 12.0 (ForteBio). Binding kinetics was evaluated using a 1:1 Langmuir model except for dimeric ACE2 and antibodies G32B6 and C12A2, which were analyzed by a bivalent binding model. Sensorgrams showing unrealistic off-rates were fit individually to a single exponential function [R=Req*(1-e^(-k*t))] to obtain the steady state response Req at each concentration. The KD was obtained by fitting Req value and its corresponding concentration to the model: “one site-specific” using GraphPad Prism 8.0.2 according to H.J. Motulsky, Prism 5 Statistics Guide, 2007, GraphPad Software Inc., San Diego CA, www.graphpad.com). All K_D_ values for multivalent interactions with antibody IgG or dimeric ACE2 and trimeric S protein are the apparent affinities with avidity effects.

### Flow cytometry

Expi293F cells (ThermoFisher Scientific) were grown in Expi293 expression medium (ThermoFisher Scientific). Cell surface display DNA constructs for the SARS-CoV-2 spike variants together with a plasmid expressing blue fluorescent protein (BFP) were transiently transfected into Expi293F cells using ExpiFectamine 293 reagent (ThermoFisher Scientific) per manufacturer’s instruction. Two days after transfection, the cells were stained with primary antibodies at 10 μg/ml concentration. For antibody staining, an Alexa Fluor 647 conjugated donkey anti-human IgG Fc F(ab’)2 fragment (Jackson ImmunoResearch, West Grove, PA) was used as secondary antibody at 5 μg/ml concentration. Cells were run through an Intellicyt iQue Screener Plus flow cytometer. Cells gated for positive BFP expression were analyzed for antibody and ACE2_615_-foldon T27W binding. The flow cytometry assays were repeated three times with essentially identical results.

### MLV-based pseudovirus assay

Murine Leukemia Virus (MLV) particles (plasmids of the MLV components kindly provided by Dr. Gary Whittaker at Cornell University and Drs. Catherine Chen and Wei Zheng at National Center for Advancing Translational Sciences, National Institutes of Health), pseudotyped with various SARS-CoV-2 S protein constructs, were generated in HEK 293T cells, following a protocol described previously for SARS-CoV (*59, 60*). To enhance incorporation of S protein into the particles, the C-terminal 19 residues in the cytoplasmic tail of each S protein were deleted. To increase the cleavage between S1 and S2, 1.5 μg of the furin expression construct was added into the DNA mixture (20 μg) for MLV particle production. To prepare for infection, 7.5×10^3^ of HEK 293 cells, stably transfected with a full-length human ACE2 expression construct, in 15 μl culture medium were plated into a 384-well white-clear plate coated with poly-D-Lysine to enhance the cell attachment. On day 2, 15 μl of MLV pseudoviruses for each variant were added into each well pre-seeded with HEK293-ACE2 cells. The plate was centrifuged at 114 xg for 5 min at 12°C. After incubation of the pseudoviruses with the cells for a time period (10 min-8 hr), as indicated in the figures, the medium was removed and the cells were washed once with 1xDPBS. 30 μl of fresh medium was added back into each well. The cells were then incubated at 37°C for additional 40 hr. Luciferase activities were measured with Firefly Luciferase Assay Kit (CB-80552-010, Codex BioSolutions Inc).

### HIV-based pseudovirus assay

Neutralizing activity against SARS-CoV-2 pseudovirus was measured using a single-round infection assay in 293T/ACE2 target cells. Pseudotyped virus particles were produced in 293T/17 cells (ATCC) by co-transfection of plasmids encoding codon-optimized SARS-CoV-2 full-length S constructs, packaging plasmid pCMV DR8.2, and luciferase reporter plasmid pHR’ CMV-Luc. G614 S, Omicron S, packaging and luciferase plasmids were kindly provided by Drs. Barney Graham and Tongqing Zhou (Vaccine Research Center, NIH). The 293T cell line stably overexpressing the human ACE2 cell surface receptor protein was kindly provided by Drs. Michael Farzan and Huihui Ma (The Scripps Research Institute). For neutralization assays, serial dilutions of monoclonal antibodies (mAbs) were performed in duplicate followed by addition of pseudovirus. Pooled serum samples from convalescent COVID-19 patients or pre-pandemic normal healthy serum (NHS) were used as positive and negative controls, respectively. Plates were incubated for 1 hour at 37°C followed by addition of 293/ACE2 target cells (1×10^4^/well). Wells containing cells + pseudovirus (without sample) or cells alone acted as positive and negative infection controls, respectively. Assays were harvested on day 3 using Promega BrightGlo luciferase reagent and luminescence detected with a Promega GloMax luminometer. Titers are reported as the concentration of mAb that inhibited 50% or 80% virus infection (IC50 and IC80 titers, respectively). All neutralization experiments were repeated twice with similar results.

### SARS-CoV-2 clinical isolates

The SARS-CoV-2 clinical isolates including an early D614G strain – New York-PV09158/2020 (NY (614G); hCoV-19/USA/NY-PV09158/2020, GenBank: MT371034 and GISAID: EPI_ISL_422525), and variants of concern (VOC) representing Delta (B.1.617.2; hCoV-19/USA/KY-CDC-2-4242084/2021, GenBank: MZ082533.1 and GISAID EPI_ISL_1823618), Omicron (B.1.1.529) BA.1 (hCoV-19/USA/CA-CDC-4358237-001/2021, GenBank: OM264909.1) and BA.2 (hCoV-19/USA/MD-HP24556/2022, GenBank: ON128736.1) lineages were obtained through BEI Resources (Manassas, VA) or Centers for Disease Control and Prevention (Atlanta, GA). Seed viruses were amplified in Vero E6 (ATCC CRL-1586) or Vero E6 with TMPRSS2 overexpression (BPS Bioscience #78081). Amplified viruses were aliquoted and stored in a secured −80 °C freezer until use. Virus titers were determined using an ELISA-based 50% tissue culture infectious dose (TCID_50_) method (*33*). All work involving infectious SARS-CoV-2 viruses was performed in an FDA Animal Biosafety Level-3 (ABSL-3) laboratory equipped with advanced access control devices and by trained personnel equipped with powered air-purifying respirators.

### In vitro virus replication and focus-forming assay

Vero-E6 cells were pre-seeded in 12-well tissue culture plates overnight and were infected with authentic viruses (G614, Delta, Omicron BA.1 or BA.2) at MOI of 0.01 in Gibco™ high glucose DMEM containing 3% FBS. Supernatants were sampled at different time points for viral titers by focus-forming assay (*61*) with some modifications. Briefly, 10-fold serially diluted postinfection were added at 100 μl/well to Vero E6-TMPRSS2 cells pre-seeded in 96-well tissue culture plates. After 1 h incubation at 37°C, 5% CO_2_, unbounded viruses were removed. Cells were overlaid with 100 μl/well of the 1:1 (v/v) mixture of 2.4% Avicel (DuPont) and 2XEMEM (Lonza) with 4% FBS. After incubation at 37°C, 5% CO_2_ for 24 hours without disturbance, plates were fixed with 10% neutral buffered formalin at room temperature for 20 min followed by permeabilization with 0.1% Triton X-100 for another 10 min. After washing, plates were probed with 1 μg/ml of inhouse developed rabbit polyclonal antibody specific for SARS-CoV-2 membrane/nucleocapsid (*33*) at 4°C overnight followed by peroxidase-conjugated goat anti-rabbit secondary antibody (SeraCare #5220-0336, 1:2000) for 2h at room temperature. Color was developed using KPL TrueBlue substrate (SeraCare #5510-0030). Foci were imaged and counted using AID vSpot Spectrum (Autoimmun Diagnostika GmbH, Strassberg, Germany) and were expressed as focus-forming units (FFU) per ml.

### Mouse study

Hemizygous B6.Cg-Tg(K18-ACE2)2Prlmn/J (K18-hACE2) transgenic mice (JAX Stock No. 034860; ref (*32*)) were bred at FDA White Oak Vivarium. All K18-hACE2 mice were genotype-confirmed (Transnetyx) and were used at 8-12 weeks old for experiments. In the ABSL-3 lab, K18-hACE2 mice were randomly grouped and were inoculated intranasally with NY (G614), Delta, Omicron BA.1 or Omicron BA.2 variant at 100 TCID_50_/mouse under isoflurane anesthesia. At 1 and 3 days post infection (dpi), infected mice were euthanized and various tissues (brain, heart, nasal turbinates, whole lung, liver, spleen and kidneys) were harvested for viral load determination. All procedures were performed according to the animal study protocols approved by the FDA White Oak Animal Program Animal Care and Use Committee.

### hACE2 expression and viral load in mouse tissues

RNeasy Plus Mini Kit (Qiagen #74136) was used to extract total RNA from mouse tissues. Mouse tissues were homogenized in sterile PBS (pH7.2) (10%, v/w) using a Fisherbrand™ Bead Mill 24 Homogenizer (FisherScientific), except nasal turbinates were directly collected in 350 μl/tube of RLT plus buffer. Extracted RNA was converted to cDNA using the High-Capacity cDNA Reverse Transcription Kit (Thermo Fisher Scientific #4368813). hACE2 expression or SARS-CoV-2 nucleocapsid (N) gene in individual mouse organs was determined using QuantiNova SYBR Green PCR kit (Qiagen #208052) in combination of 500 nM of hACE2 gene specific primer set (Integrated DNA Technologies, Assay ID: Hs.PT.58.27645939) or 500 nM of 2019-nCoV RUO Kit (Integrated DNA Technologies #10006713), respectively. The cycling program was performed in Stratagene MX3000p qPCR system (Agilent) as follows: 95 °C for 120 s, 95 °C for 5 s (50 cycles) and 60 °C for 18 s (*33*). Serially diluted pCMV6-AC-ACE2-GFP plasmid or pCC1-CoV2-F7 plasmid expressing SARS-CoV-2 N (*62*) was used to construct a standard curve. The copies of expressed hACE2 or SARS-CoV-2 N gene copies in individual tissues were interpolated based on threshold cycle (Ct) values determined using MxPro qPCR software (Agilent). A value of 1 was assigned if gene copies were below the detection limits.

### Cryo-EM sample preparation and data collection

To prepare cryo EM grids, the purified full-length BA.2 spike protein at 4.6 mg/ml in DDM was applied to a 1.2/1.3 Quantifoil gold grid (Quantifoil Micro Tools GmbH), which had been glow discharged with a PELCO easiGlow™ Glow Discharge Cleaning system (Ted Pella, Inc.) for 60 s at 15 mA. Grids were immediately plunge-frozen in liquid ethane using a Vitrobot Mark IV (ThermoFisher Scientific), and excess protein was blotted away using grade 595 filter paper (Ted Pella, Inc.) with a blotting time of 4 s, a blotting force of −12 at 4°C with 100% humidity. The grids were first screened for ice thickness and particle distribution. Selected grids were used to acquire images with a Titan Krios transmission electron microscope (ThermoFisher Scientific) operated at 300 keV and equipped with a BioQuantum GIF/K3 direct electron detector. Automated data collection was carried out using SerialEM version 3.8.6 (*63*) at a nominal magnification of 105,000× and the K3 detector in counting mode (calibrated pixel size, 0.83 Å) at an exposure rate of 13.761 electrons per pixel per second. Each movie add a total accumulated electron exposure of ~53.853 e-/Å^2^, fractionated in 50 frames. Data sets were acquired using a defocus range of 0.5-2.2 μm.

### Image processing and 3D reconstructions

All data were processed using cryoSPARC v.3.3.1 (*36*). Drift correction for cryo-EM images was performed using patch mode, and contrast transfer function (CTF) estimated by patch mode. Motion corrected sums with dose-weighting were used for all other image processing. Templates were first generated using manually picked particles and template-based particle picking was then performed. 6,763,049 particles were extracted from 34,833 images using a box size of 672Å (downsizing to 128Å). The particles were subjected to four rounds of 2D classification, giving 2,749,398 good particles. A low-resolution negative-stain reconstruction of the Wuhan-Hu-1 (D614) sample was low-pass filtered to 40Å resolution and used as an initial model. The selected good particles were first used for one round of heterogeneous classification with six copies of the initial model as the reference in C1 symmetry. A major class (21.1%) with clear structural features was re-extracted to a smaller box size (480Å) and subjected to another round of heterogeneous refinement with four copies of the initial model as the reference in C1 symmetry. Two major classes, representing two different one-RBD-up conformations, were combined and subjected to another round of heterogeneous refinement with four copies of the initial model as the reference in C1 symmetry, giving four major classes, one for a three-RBD-down conformation, two for a one-RBD-up conformation and one for a one-RBD-intermediate conformation. The two one-RBD-up classes were combined and subjected to one round of non-uniform refinement in C1 symmetry, giving a map at 3.1Å resolution from 193,655 particles. One more round of local CTF refinement was performed to further improve the resolution to 3.0Å. The two three-RBD-down classes from the second round of heterogeneous refinement were combined with the three-RBD-down class and one-RBD-intermediate class from the third round of heterogeneous refinement, and subjected to one additional round of heterogeneous refinement with six copies of the initial model as the reference in C1 symmetry. There were two major classes, representing the three-RBD-down conformation and one-RBD-intermediate conformation, respectively. These two classes were further refined in C3 symmetry for the three-RBD-down conformation, giving a map at 3.0Å resolution from 160,774 particles; and in C1 symmetry for one-RBD-intermediate conformation, giving a map at 3.3Å resolution from 113,667 particles. They were subjected to local CTF refinement to improve the overall resolution, resulting in a 2.8Å map and a 3.1Å map, respectively. To improve the resolution in the RBD and NTD, several different local refinements were performed with a soft mask covering both the RBD and NTD regions. For the three-RBD-down conformation, the local refinement gave a map at 3.0Å resolution. Likewise, the local refinement gave a 3.3Å map for the one-RBD-up conformation, and a 3.5Å map for the one-RBD-intermediate conformation. The density near the furin cleavage site in the cryo-SPARC maps was weak probably due to insufficient local classification. We therefore transferred the selected particles from cryoSPARC to RELION (*64*), and performed local classification near the cleavage site without alignment. Each conformation gave a major class, which was subjected to another round of 3D autorefinement. The RELION maps showed strong density for the extra residues first observed for the Omicron BA.1 subvariant (*29*), but not for other variants (*26–28, 30*). The best density maps were used for model building.

All resolutions were reported from the gold-standard Fourier shell correlation (FSC) using the 0.143 criterion. Density maps were corrected from the modulation transfer function of the K3 detector and sharpened by applying a temperature factor that was estimated using Sharpening Tools in cryoSPARC. Local resolution was also determined using cryoSPARC.

### Model building

The initial templates for model building were our G614 S trimer structures (PDB ID: 7KRQ and PDB ID: 7KRR; ref(*26*)). Several rounds of manual building were performed in Coot. The model was then refined in Phenix (*65*) against the 2.8Å (three-RBD-down conformation), 3.0Å (one-RBD-up conformation), 3.1Å(one-RBD-intermediate conformation) cryo-EM maps of the BA2 S trimer. Iteratively, refinement was performed in both Phenix (real space refinement) and ISOLDE (*66*), and the Phenix refinement strategy included minimization_global, local_grid_search, and adp, with rotamer, Ramachandran, and reference-model restraints, using 7KRQ and 7KRR as the reference models. The N-terminal segment of the is shortened by the three-residue deletion (L24del-P25del-P26del) and also constrained by the disulfide bond between Cys15 and Cys136. There is reasonable density in which the 70-80 loop, disordered in many previous S trimer structures, could be modeled (Fig. S16). Such a structured loop can create a knot in this region, however, which will need a higher resolution map to confirm. The refinement statistics are summarized in Table S3. Structural biology applications used in this project were compiled and configured by SBGrid (*67*).

## Acknowledgments

We thank the SBGrid team for computing support, computing resources from S. Harrison and J. Abraham, K. Arnett for support and advice on the BLI experiments, and S. Harrison for critical reading of the manuscript. EM data were collected at the Harvard Cryo-EM Center for Structural Biology of Harvard Medical School. We acknowledge support for COVID-19 related structural biology research at Harvard from the Nancy Lurie Marks Family Foundation and the Massachusetts Consortium on Pathogen Readiness (MassCPR). This work was supported by Fast grants by Emergent Ventures (to B.C. and D.R.W.), COVID-19 Awards by MassCPR (to B.C., D.R.W., and M.S.S.), and NIH grants AI147884 (to B.C.), AI141002 (to B.C.), AI127193 (to B.C. and James Chou) and AI39538, AI165072, and AI169619 (to D.R.W). This work was also supported in part by FDA/CBER intramural SARS-CoV-2 pandemic fund (to H.X.). The clinical isolate New York-PV09158/2020 (ATCC #NR-53516) and Omicron (B.1.1.529) BA.1 were obtained through BEI Resources, NIAID, NIH: SARS-Related Coronavirus 2. The Delta (B. 1.617.2) and Omicron BA.2 seed viruses were kindly provided by Drs. Bin Zhou and Charles (Todd) Davis at Centers for Disease Control and Prevention, Atlanta, GA.

## Author Contribution

B.C., T.X., J.Z. and H.G. conceived the project. H.G. expressed and purified the full-length S proteins with help from H.P.. T.X. designed and performed BLI and cell-cell fusion experiments, also assisted by H.G.. J.Z. prepared cryo grids and performed EM data collection with contributions from M.L.M., and processed the cryo-EM data, built and refined the atomic models. W.T., M.K., H.J.K. and H.X. carried out the animal study and in vitro virus replication kinetics using authentic viruses. C.L.L. and M.S.S performed the neutralization assays using the HIV-based pseudoviruses. J.L. and S.W. created the BA.2 expression construct and performed the neutralization assays using the MLV-based pseudoviruses. H.Z., K.A. and W.Y. performed the flow cytometry experiments. P.T., A.G. and, D.R.W. produced anti-S monoclonal antibodies. S.R.V. contributed to cell culture and protein production. All authors analyzed the data. B.C., T.X., J.Z., H.G. and H.X. wrote the manuscript with input from all other authors.

