## Supplemental Figures and Tables for "Structural and functional characteristics of SARS-CoV-2 Omicron subvariant BA.2 spike"

### Supplementary materials

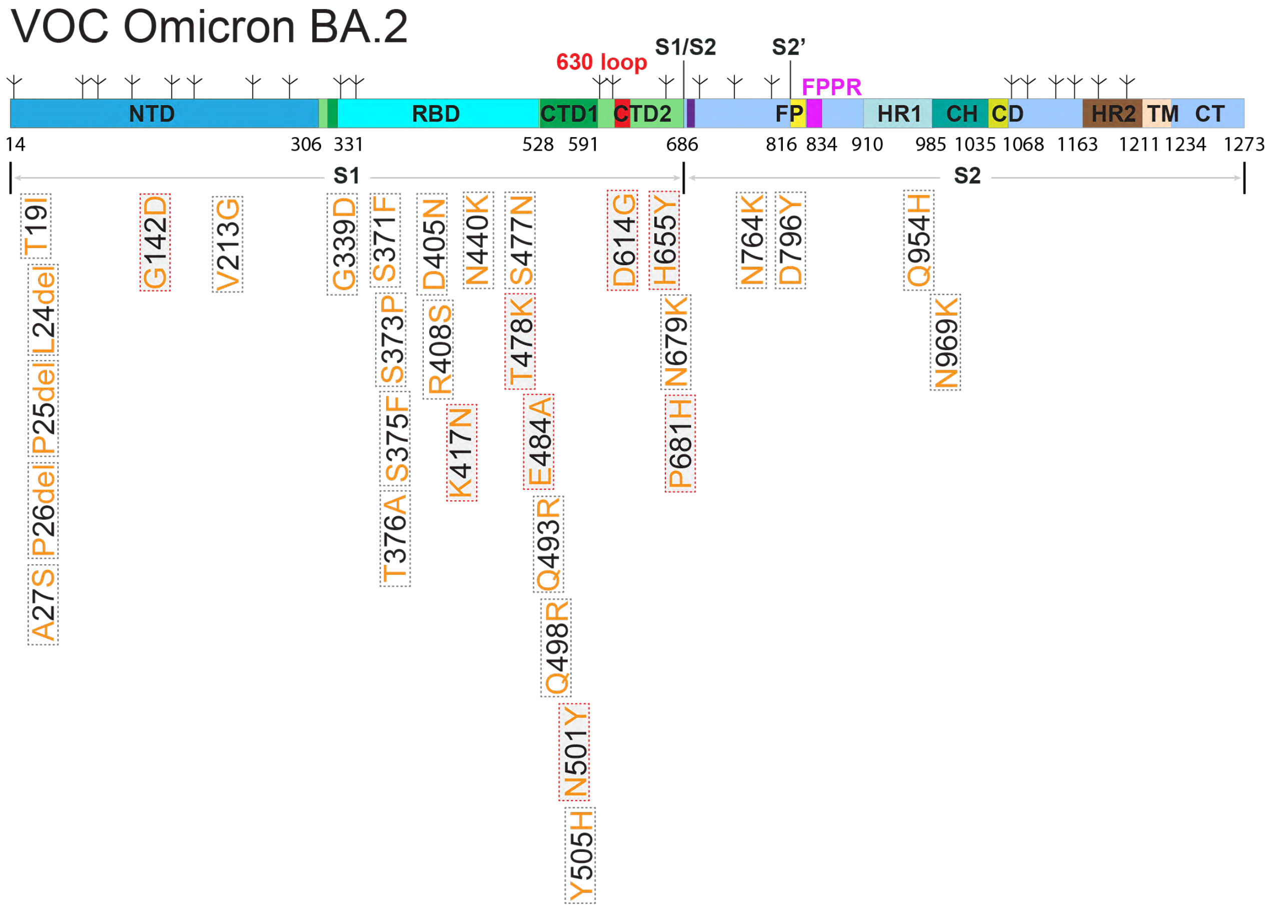

**Figure S1. Schematic representation of a full-length Omicron BA.2 spike (S) protein.** The sequence is derived from an Omicron BA.2 subvariant (hCoV-19/Denmark/DCGC-327158/2022). Segments of S1 and S2 include: NTD, N-terminal domain; RBD, receptor-binding domain; CTD1, C-terminal domain 1; CTD2, C-terminal domain 2; 630 loop, residues 620-640; S1/S2, the furin cleavage site at the S1/S2 boundary; S2', S2' cleavage site; FP, fusion peptide; FPPR, fusion peptide proximal region; HR1, heptad repeat 1; CH, central helix region; CD, connector domain; HR2, heptad repeat 2; TM, transmembrane segment; CT, cytoplasmic tail; and tree-like symbols for glycans. Positions of all mutations (from the amino-acid sequence of Wuhan-Hu-1) are indicated and those highlighted in red rectangles are also present in at least one of the previous VOCs.

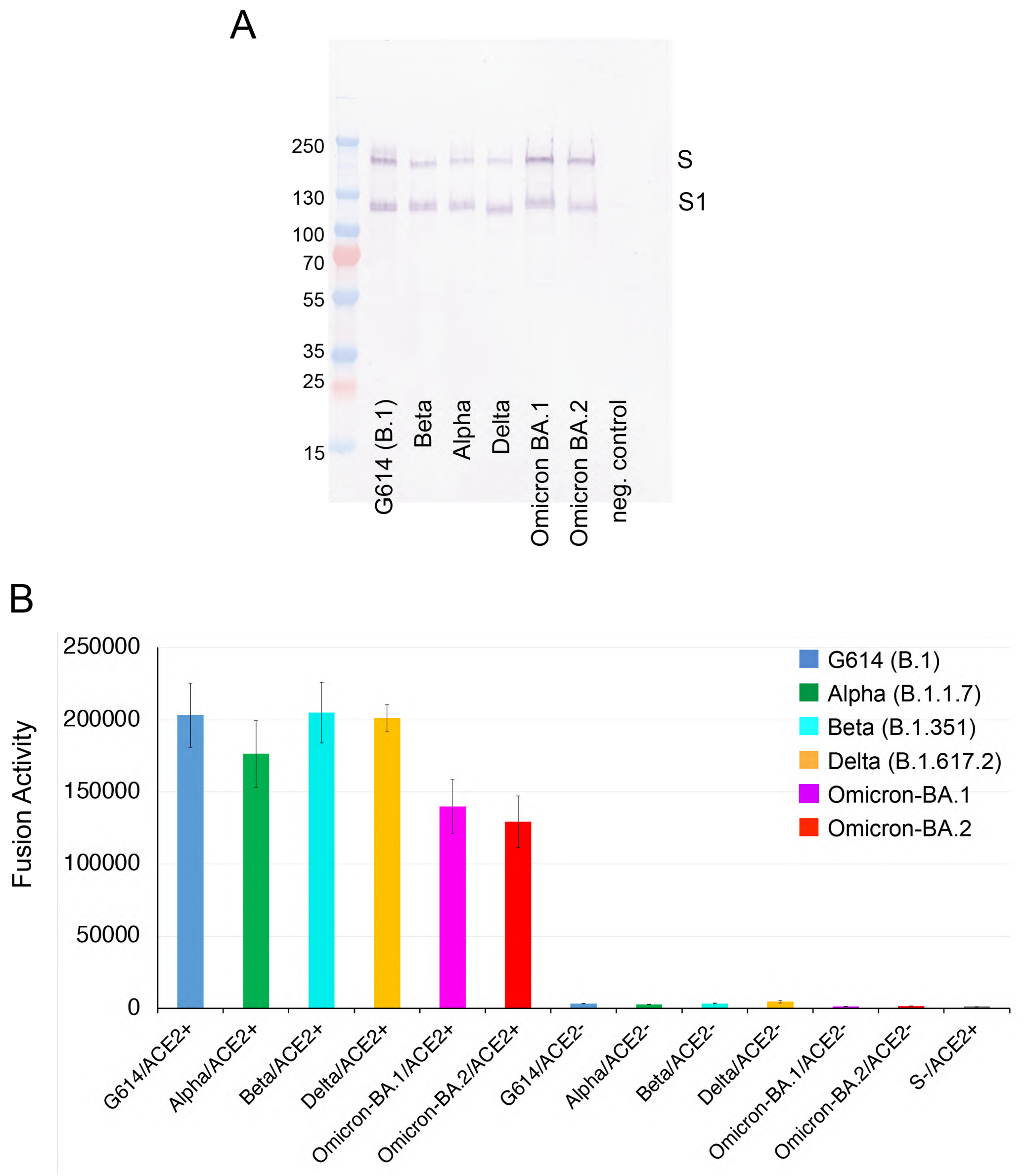

**Figure S2. Expression and cell-cell fusion of the S protein from Omicron BA.2.** (**A**) Expression and processing of the full-length S constructs of various variants in HEK293 cells. S protein samples prepared from HEK293 cells transiently transfected with 10 μg of the full-length S expression plasmids were detected by anti-RBD polyclonal antibodies. Bands for the uncleaved S and S1 fragment are indicated. (**B**) HEK293T cells transfected with the untagged, full-length S protein expression plasmids were fused with ACE2-expressing cells. Cell-cell fusion led to reconstitution of α and ω fragments of β-galactosidase to form an active enzyme, and the fusion activity was then quantified by a chemiluminescent assay. No ACE2 and no S were negative controls.

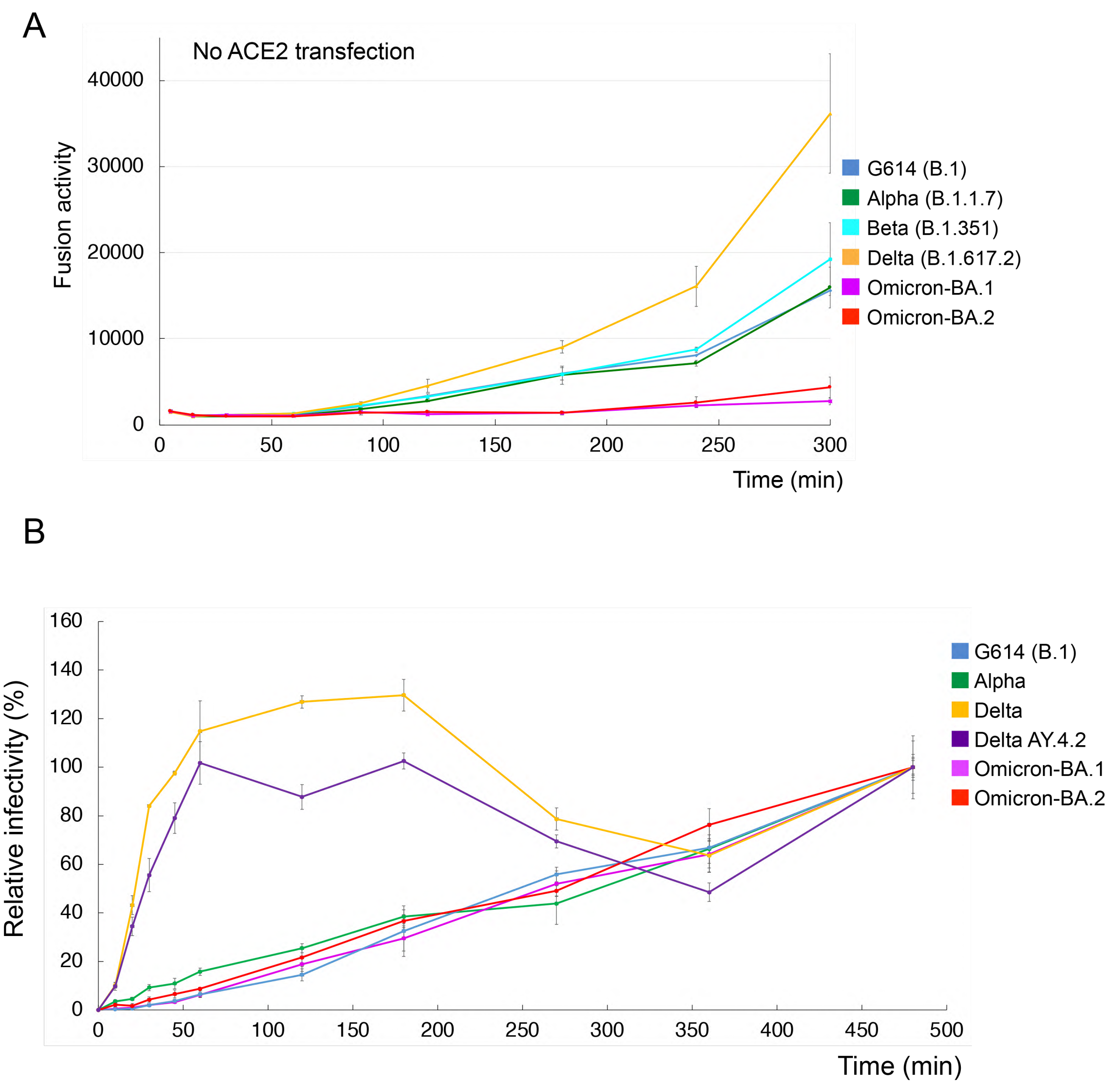

**Figure S3. Infection of HEK293-ACE2 cells by MLV-based pseudotyped viruses.** (**A**) Time-course of cell-cell fusion mediated by various full-length S proteins, as indicated, using HEK293 cells with endogenous ACE2 only. (**B**) Time course for single-cycle infection of HEK293-ACE2 cells by MLV-based pseudotyped viruses with various SARS-CoV-2 variant S constructs, as indicated, all containing a CT deletion. Infection was initiated by mixing viruses and target cells, and viruses were washed out at each time point as indicated. Delta AY.4.2 is a Delta subvariant. The experiments were repeated at least three times with independent samples giving similar results.

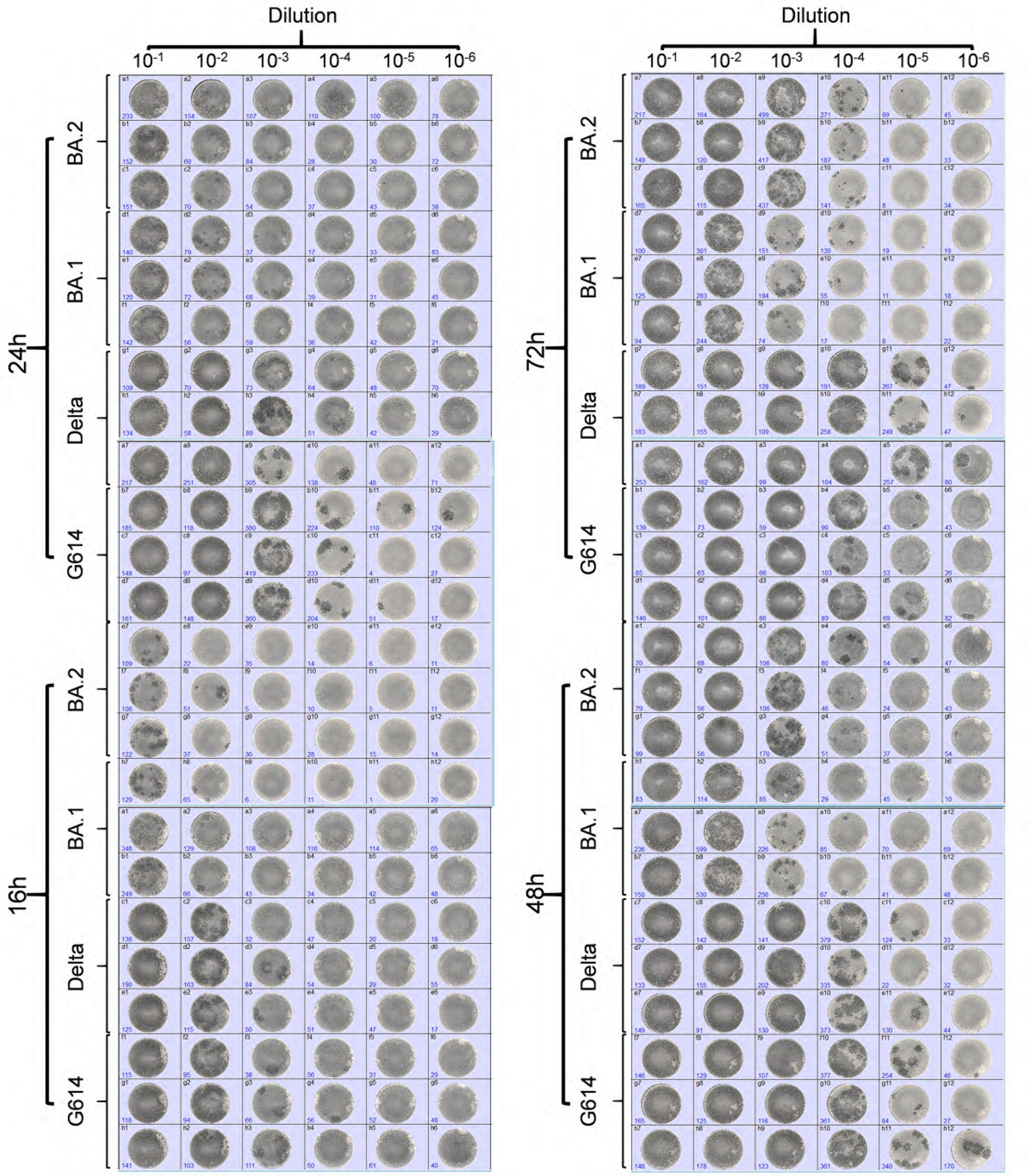

**Figure S4. Foci images of in vitro replication kinetics of the authentic viruses.** Vero E6 cells were infected with the authentic G614 (B.1), Delta (B.1617.2), Omicron-BA.1 or BA.2 viruses at MOI of 0.01. Postinfection supernatants were titrated by a focus-forming assay. Foci as a cluster of cells expressing viral antigen were imaged and counted using AID vSpot Spectrum. The number of foci (dark spots) counted is shown in the lower corner of each well.

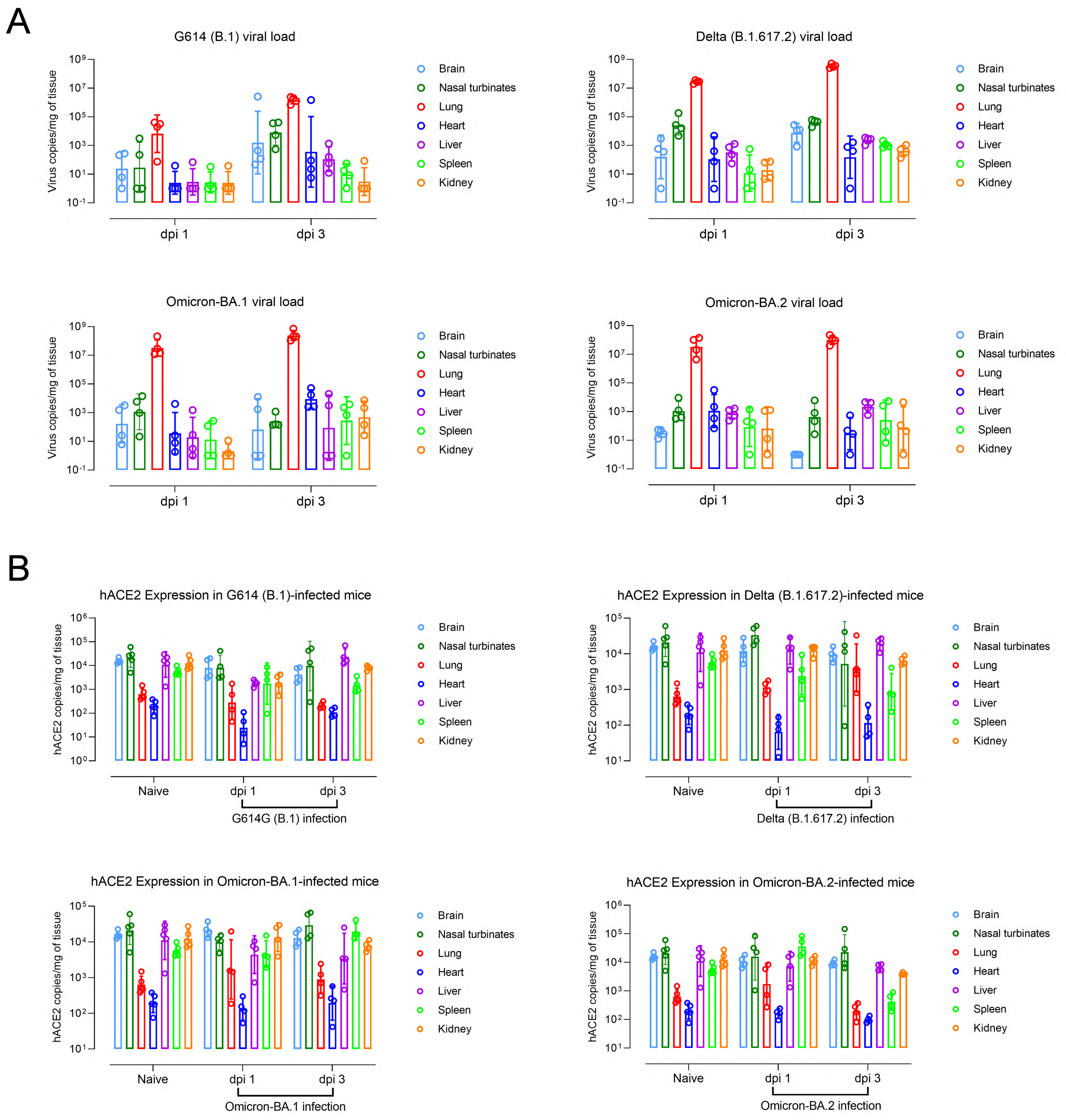

**Figure S5. Tissue-specific viral burdens and hACE2 expression in K18-hACE2 mice at 1 and 3 days post infection (dpi).** Mice were intranasally inoculated with 100 $TCID_{50}$/mouse of G614 (B.1), Delta (B.1617.2), Omicron-BA.1 or BA.2. The viral RNA copies (A) or hACE2 expression (B) in various tissue homogenates were measured by RT-qPCR. Data are expressed as geometric means (bars) with geometric standard deviation (error bars). Individual results of infected mice (n=4 mice/time point/group) and uninfected naïve mice (n=5) are shown.

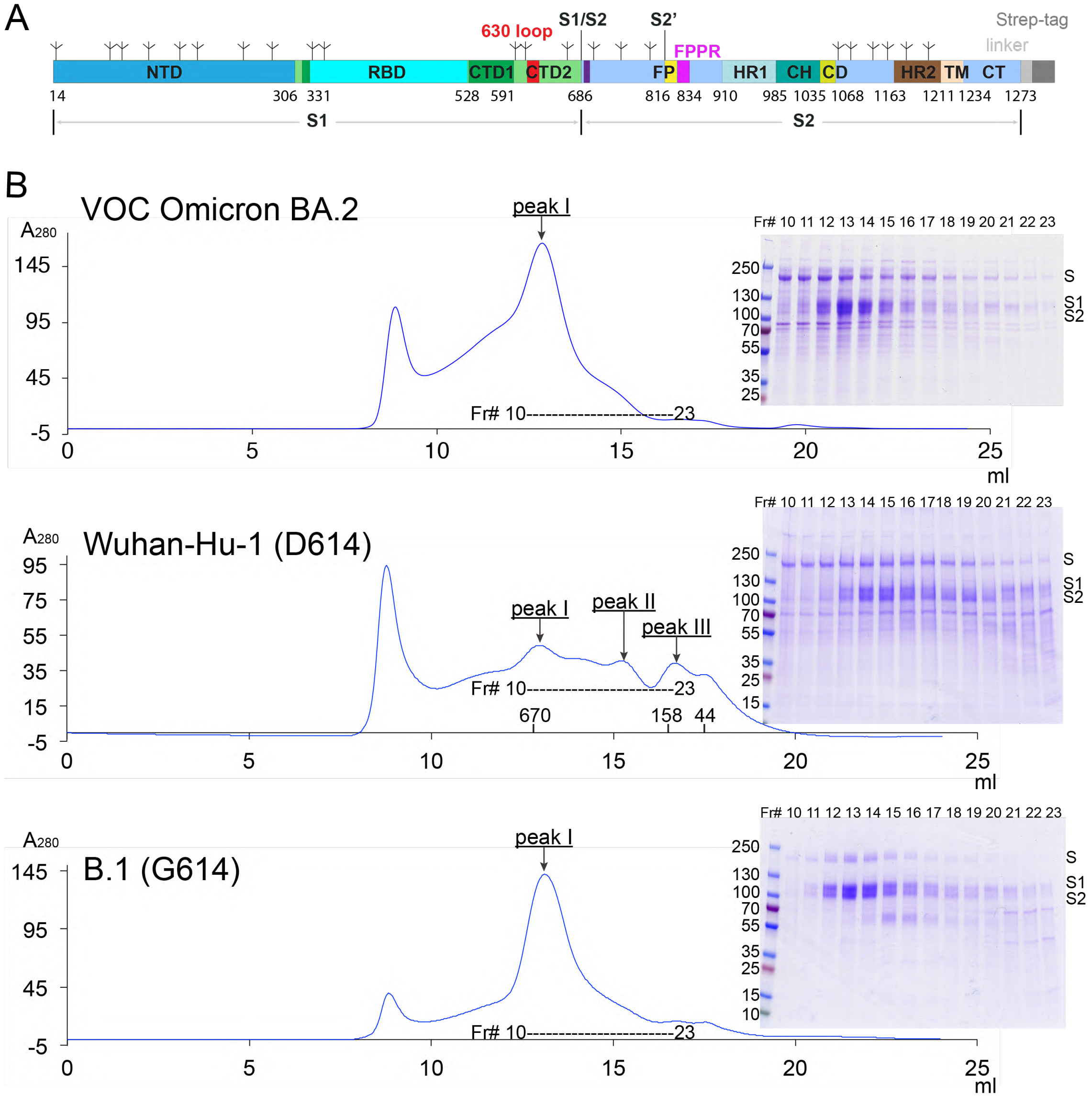

**Figure S6. Purification of the Omicron BA.2 full-length S protein.** (**A**) A strep-tag for purification was fused to the C-terminus of the full-length S protein by a flexible linker. (**B**) The full-length BA.2 S protein was extracted and purified in detergent DDM, and resolved by gel-filtration chromatography on a Superose 6 column. Peak I, the prefusion S trimer; peak II, the postfusion S2 trimer; and peak III, the dissociated monomeric S1. Inset, peak fractions were analyzed by Coomassie stained SDS-PAGE. Labeled bands are S, S1 and S2. Fr#, fraction number. Each experiment was repeated at least three times independently with similar results. The data for the preparations from the Wuhan-Hu-1 (D614) and B.1 (G614), published previously (*26, 30*), are included for convenient comparison.

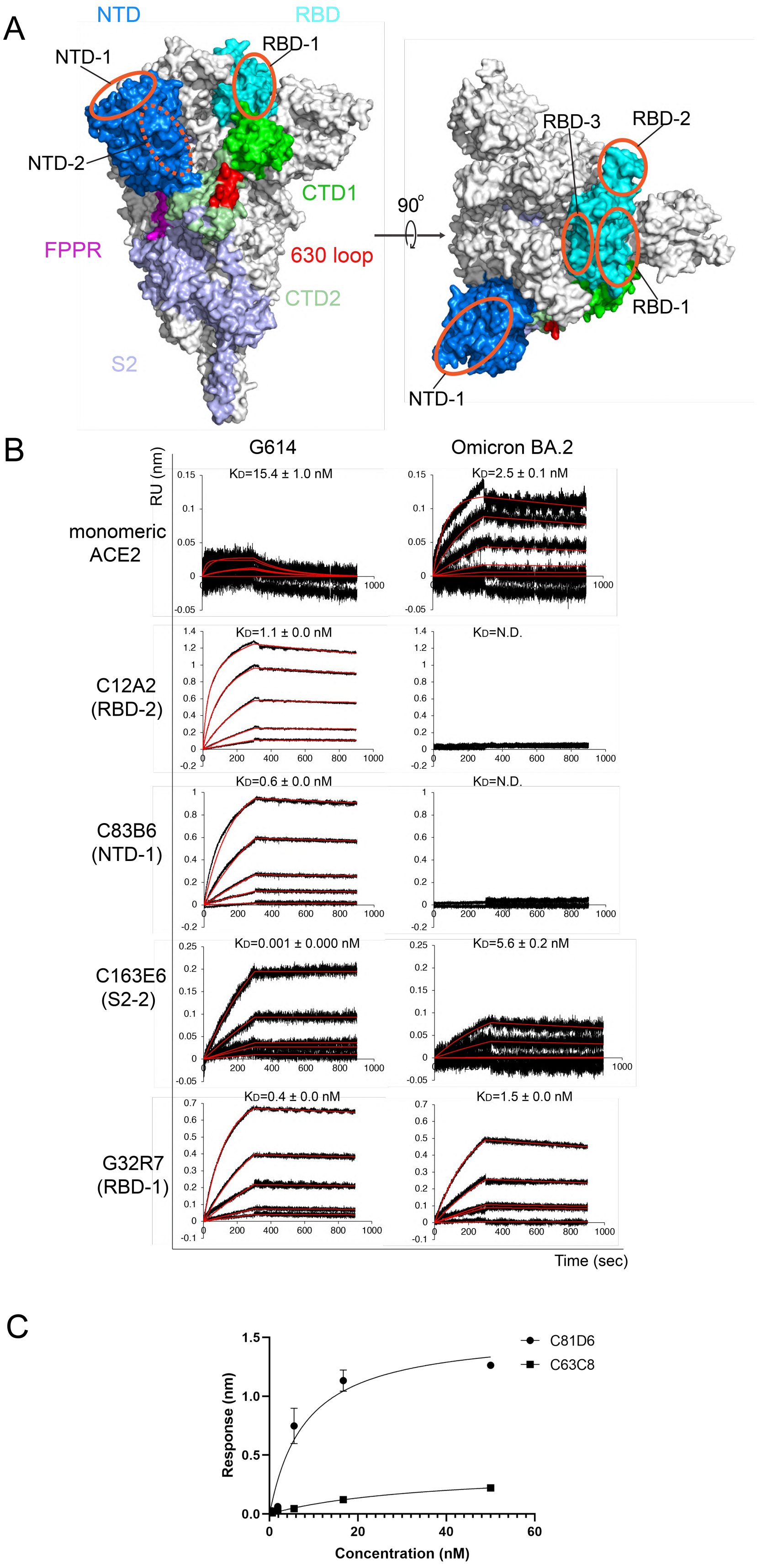
A
NTD
RBD
NTD-1
RBD-1
NTD-2
CTD1
FPPR
630 loop
CTD2
S2
90°
RBD-3
RBD-2
RBD-1
NTD-1
B
G614
Omicron BA.2
RU (nm)
monomeric ACE2
KD=15.4 ± 1.0 nM
KD=2.5 ± 0.1 nM
C12A2 (RBD-2)
KD=1.1 ± 0.0 nM
KD=N.D.
C83B6 (NTD-1)
KD=0.6 ± 0.0 nM
KD=N.D.
C163E6 (S2-2)
KD=0.001 ± 0.000 nM
KD=5.6 ± 0.2 nM
G32R7 (RBD-1)
KD=0.4 ± 0.0 nM
KD=1.5 ± 0.0 nM
Time (sec)
C
Response (nm)
Concentration (nM)
C81D6
C63C8

**Figure S7. Additional antigenic analysis of the purified full-length BA.2 S protein**. (**A**) Antibody competition groups as described in ref(*34*). Surface regions of the S trimer targeted by antibodies on S1 are highlighted by orange ellipses, including RBD-1, RBD-2, RBD-3, NTD-1 and NTD-2. The exact location of NTD-2 is uncertain and therefore marked with a dashed line. (**B**) Binding analysis of the prefusion S trimers from G614 and BA.2 with soluble monomeric ACE2 and selected monoclonal antibodies was performed by BLI. For ACE2 binding, purified ACE2 protein was immobilized to AR2G biosensors and dipped into the wells containing each purified S proteins at various concentrations. For antibody binding, various antibodies were immobilized to AHC biosensors and dipped into the wells containing each purified S protein at different concentrations. Binding kinetics were evaluated using a 1:1 Langmuir model except for antibody 12A2 targeting the RBD-2, which was analyzed by a bivalent binding model. The sensorgrams are in black and the fits in red. Binding constants highlighted by underlines were estimated by steady-state analysis as described in the Methods. RU, response unit. Binding constants are also summarized here and in Table S1. N.D., not determined. All experiments were repeated at least twice with essentially identical results. (**C**) Steady-state analysis by plotting steady-state responses against concentrations. $K_D$ values were derived from the fits.

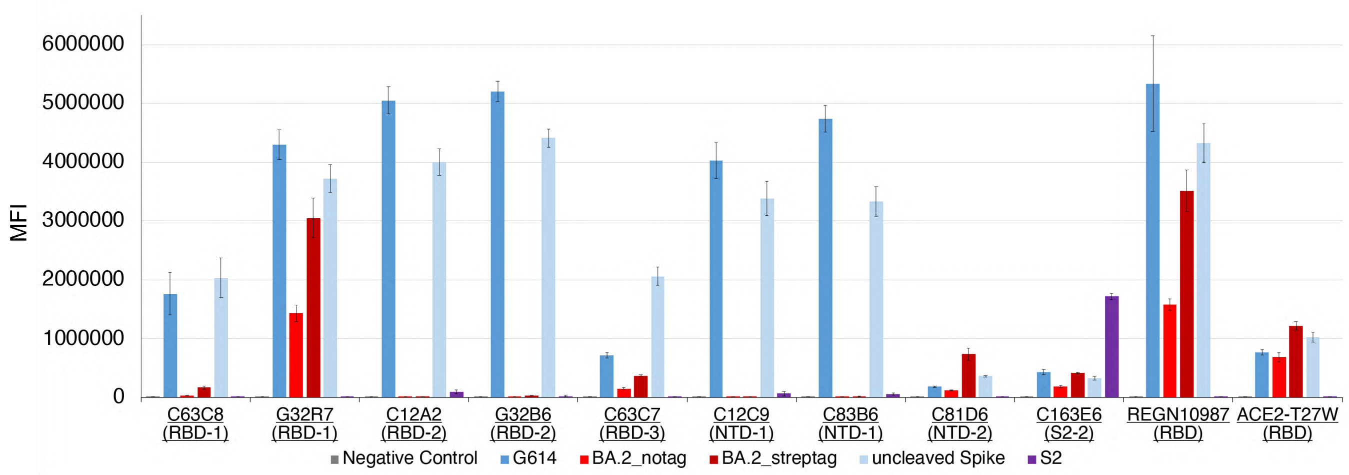

**Figure S8. Antigenic properties of the cell-surface BA.2 S protein assessed by flow cytometry.** Antibody binding to the full-length S proteins of the G614 and Omicron variants, as well as the uncleaved wildtype spike and an S2 construct expressed on the cell surfaces analyzed by flow cytometry. BA.2_notag, the unmodified, full-length S protein from the BA.2 subvariant. BA.2_streptag, the intact BA.2 S protein fused with a C-terminal twin Strep tag. The antibodies and their targets are indicated. A designed ACE2-based fusion inhibitor ACE2615-foldon-T27W was used for detecting receptor binding (*35*). MFI, mean fluorescent intensity. The error bars represent standard errors of mean from measurements using three independently transfected cell samples. The flow cytometry assays were repeated three times with essentially identical results.

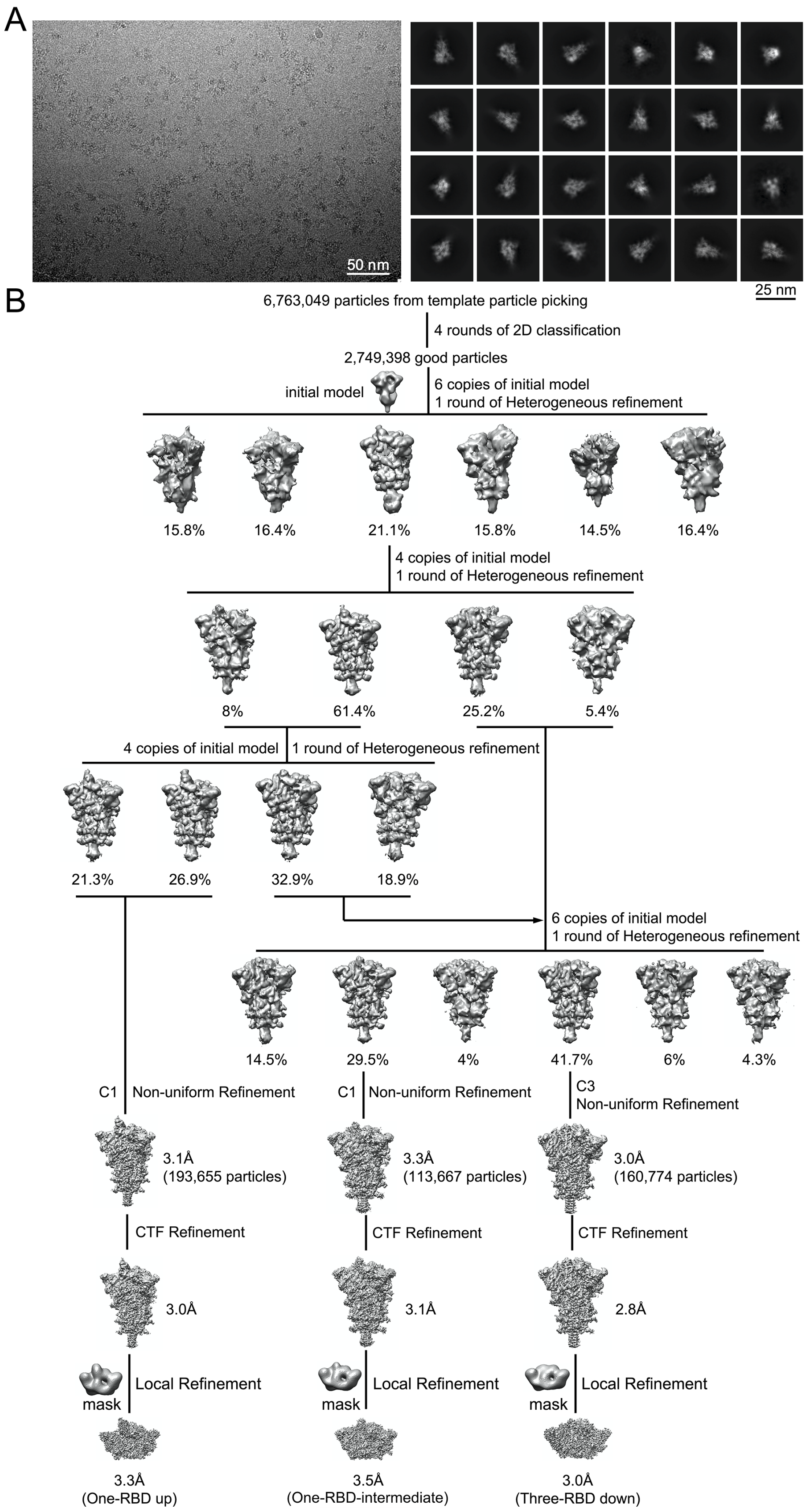
A
50 nm
25 nm
B
6,763,049 particles from template particle picking
4 rounds of 2D classification
2,749,398 good particles
initial model
6 copies of initial model
1 round of Heterogeneous refinement
15.8%
16.4%
21.1%
15.8%
14.5%
16.4%
4 copies of initial model
1 round of Heterogeneous refinement
8%
61.4%
25.2%
5.4%
4 copies of initial model
1 round of Heterogeneous refinement
21.3%
26.9%
32.9%
18.9%
6 copies of initial model
1 round of Heterogeneous refinement
14.5%
29.5%
4%
41.7%
6%
4.3%
C1
Non-uniform Refinement
C1
Non-uniform Refinement
C3
Non-uniform Refinement
3.1Å
(193,655 particles)
3.3Å
(113,667 particles)
3.0Å
(160,774 particles)
CTF Refinement
CTF Refinement
CTF Refinement
3.0Å
3.1Å
2.8Å
mask
Local Refinement
mask
Local Refinement
mask
Local Refinement
3.3Å
(One-RBD up)
3.5Å
(One-RBD-intermediate)
3.0Å
(Three-RBD down)

**Figure S9. Cryo-EM analysis of the BA.2 S trimer.** Top, representative micrograph**,** and 2D averages (box dimension: 396Å) of the cryo-EM particle images of the BA.2 S trimer. Bottom, data processing workflow for structure determination.

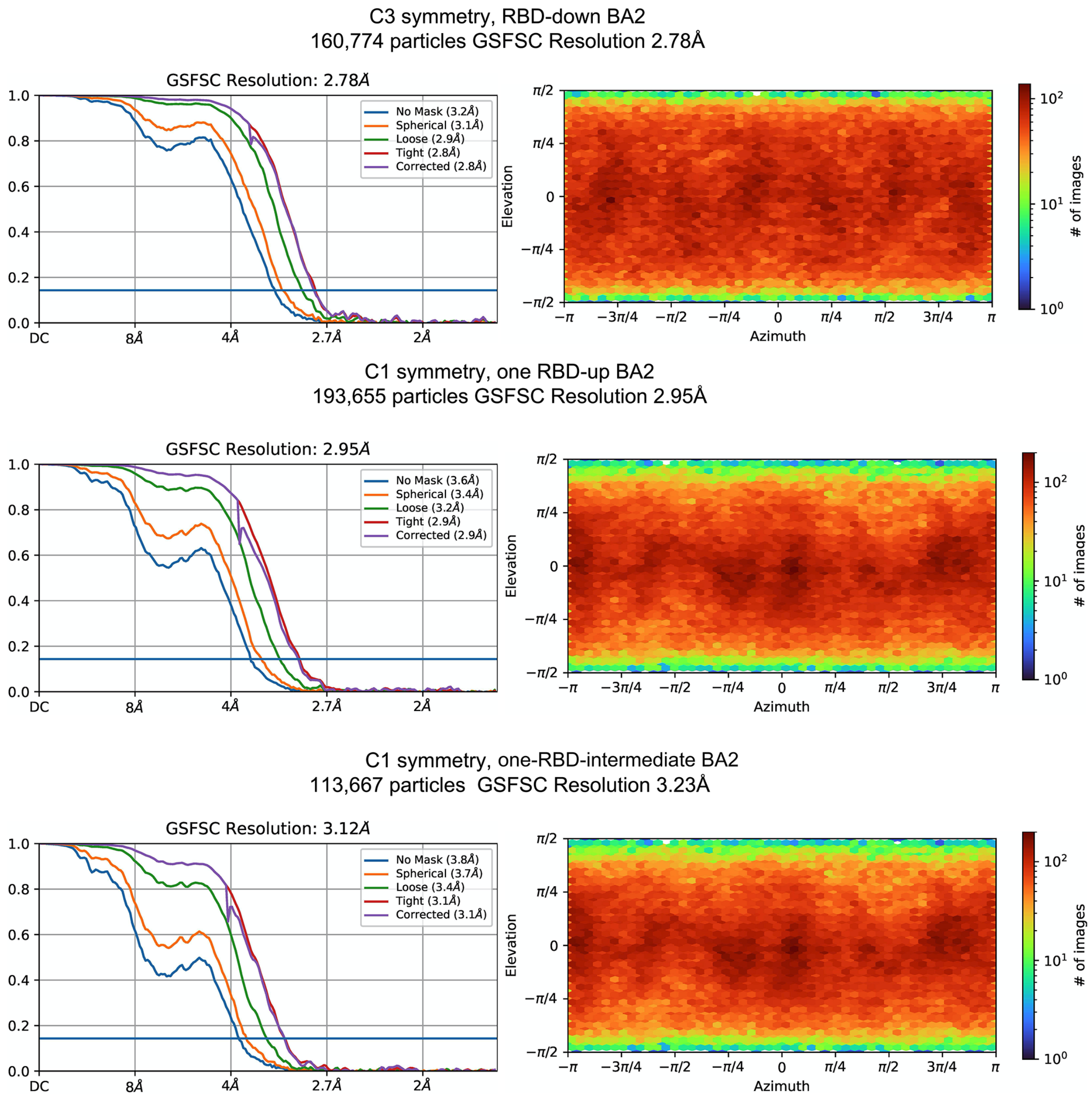

**Figure S10. Analysis of the BA.2 S trimer structures by cryoSPARC.** Gold standard FSC curves of the three refined 3D reconstructions of the BA.2 S trimer and the corresponding cryoSPARC output for particle distribution of each reconstruction.

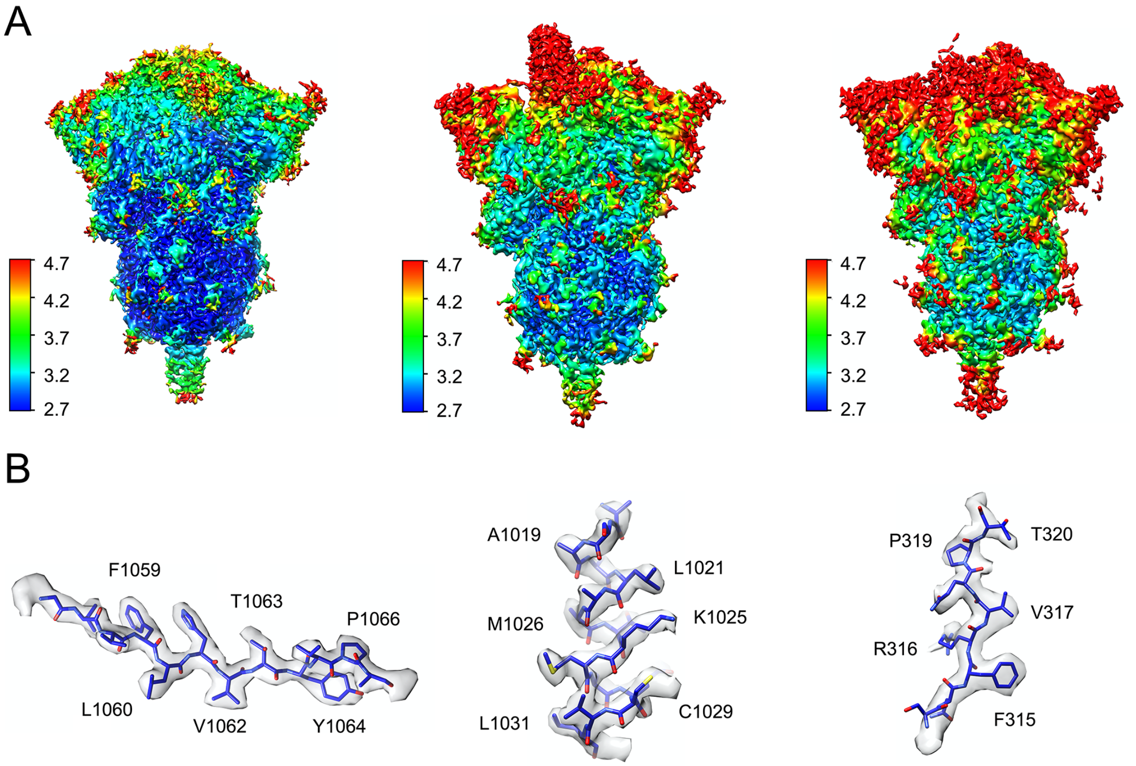

**Figure S11. Additional analysis of the BA.2 S trimer structures.** (**A**) 3D reconstructions of the BA.2 S trimer in the three-RBD-down, one-RBD-intermediate and one-RBD-up conformations, respectively, are colored according to local resolution estimated by cryoSPARC. (**B**) Representative density in gray surface from the EM map of the three-RBD-down conformation.

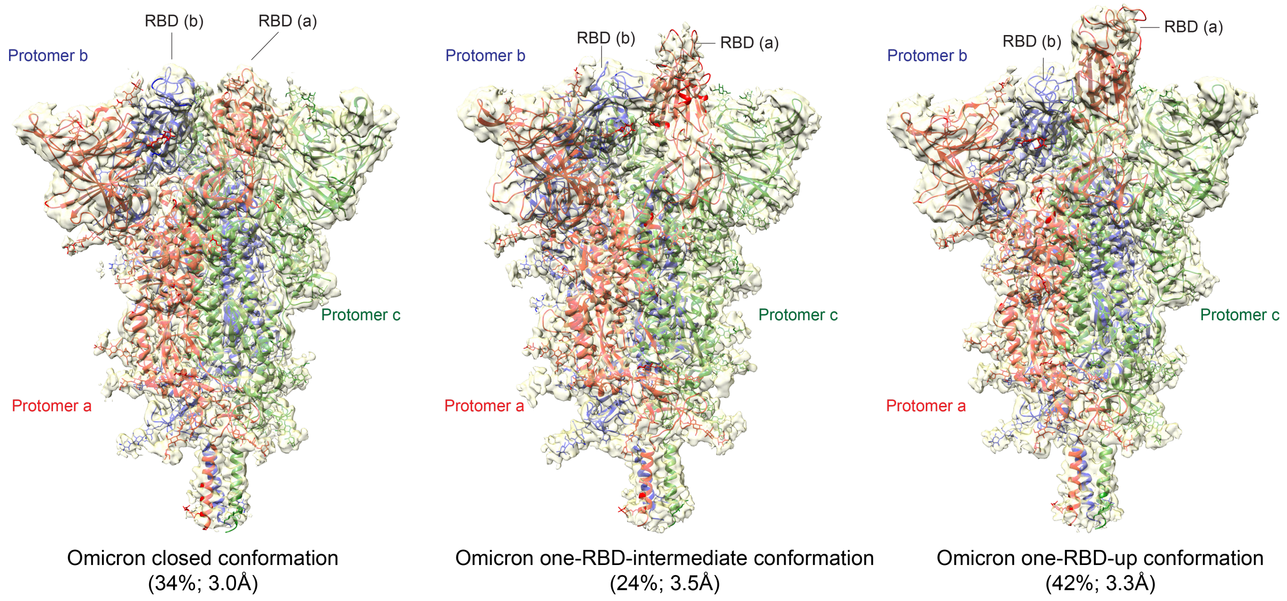

**Figure S12. Cryo-EM structures of the full-length BA.2 S protein**. Three structures of the BA.2 S trimer, representing the closed prefusion conformation, one-RBD-intermediate conformation and one-RBD-up conformations, were modeled based on corresponding cryo-EM density maps at 3.0Å, 3.5Å and 3.3Å resolution, respectively. Three protomers (a, b, c) are colored in red, blue and green, respectively. RBD locations are indicated. Particle percentage for each class in the data processing is also indicated, but it may not accurately reflect the conformation distribution of the S trimer in solution.

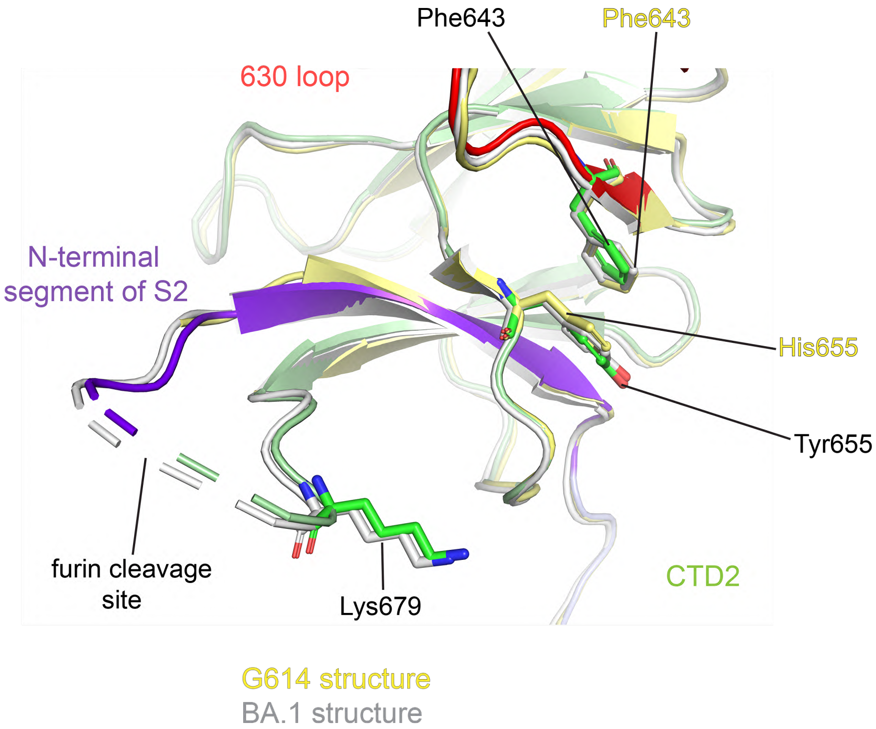

**Figure S13. Additional ordered residues near the furin site in the BA.2 structure**. Superposition of the structure of the BA.2 S trimer in ribbon representation and various colors with the structures of the BA.1 S in gray and G614 S in yellow aligned by S2, showing the region near the furin cleavage site.

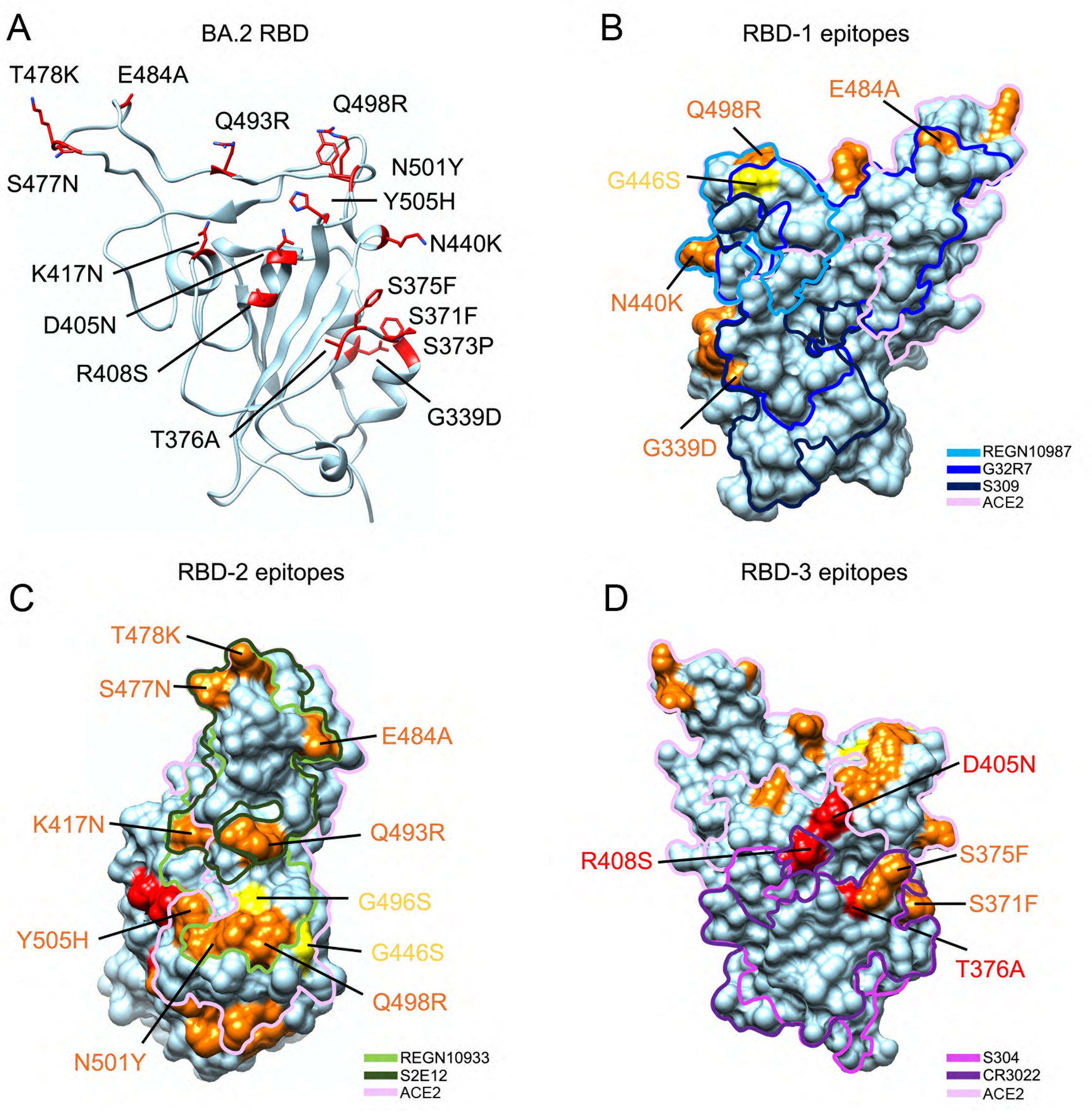

**Figure S14. Antigenic surfaces of the RBD.** (**A**) The RBD from the BA.2 S trimer structure is shown in ribbon diagram in light blue. 16 mutations are highlighted in stick model in red. (**B**) Footprints of RBD-1 antibodies, REGN10987 (*68*), G32R7 (*69*) and S309 (*69*), and ACE2 are marked on the RBD surface. The mutations shared between BA.1 and BA.2 are shown in orange; those unique to BA.1 in yellow; those unique to BA.2 in red. (**C**) Footprints of RBD-2 antibodies, REGN10933 (*68*) and S2E12 (*70*), and ACE2 are marked on the RBD surface. (**D**) footprints of RBD-3 antibodies, S304 (*71*) and CR3022 (*72*), and ACE2 on the RBD surface.

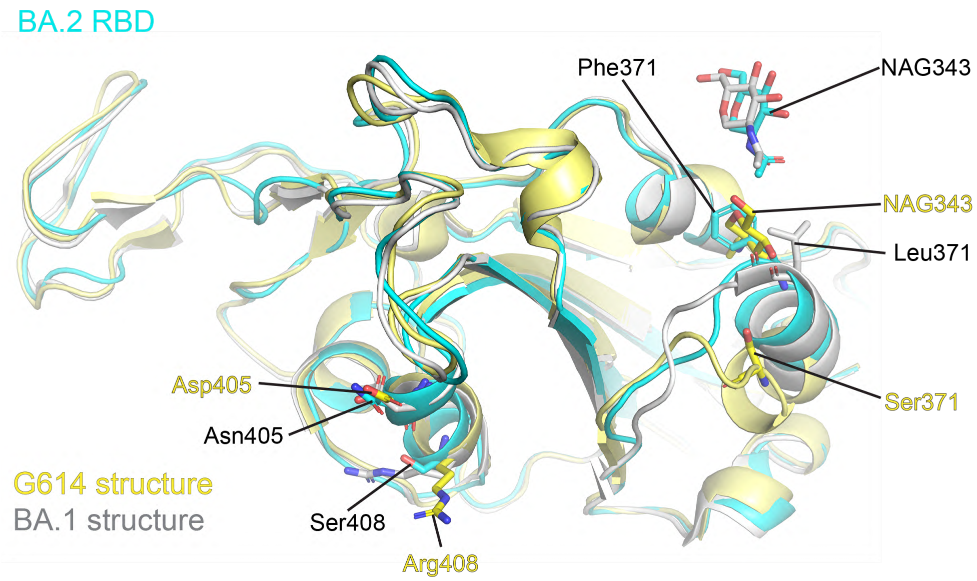

**Figure S15. A close-up view of the unique mutations in the BA.2 RBD.** Superposition of the BA.2 RBD structure in ribbon representation and cyan with the structures of the RBDs of G614 S in yellow and BA.1 in gray. The mutated residues and the N-linked glycans at Asn343 are in stick model. NAG, N-acetylglucosamine.

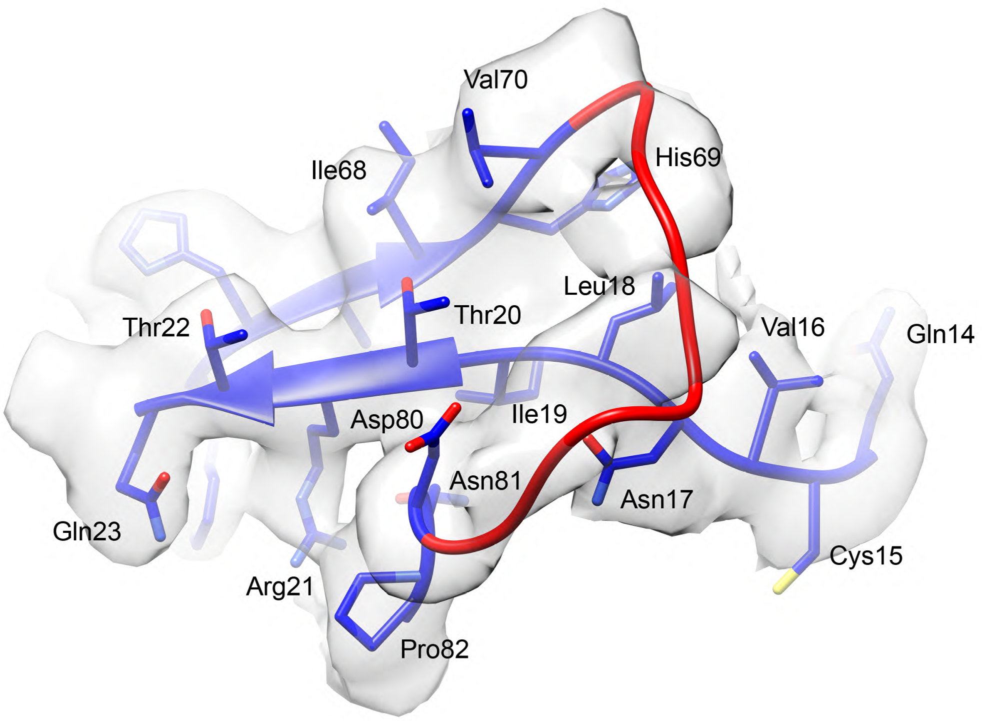

**Figure S16. Possibly ordered 70-80 loop.** The N-terminal segment of the BA.2 S is shortened by the three-residue deletion (L24del-P25del-P26del) and also constrained by the disulfide bond between Cys15 and Cys136. There is reasonable density in which the 70-80 loop, disordered in many previous S trimer structures, could be modeled (shown in red). Such a structured loop can create a knot in this region, however, which will need a higher resolution map to confirm.

### Table S1. Binding constants of S-ACE2/antibody interactions

| | | $K_D$ (M) | $K_D$ Error | ka (1/Ms) | ka2 | ka Error | ka2 Error | kdis (1/s) | kdis2 | kdis Error | kdis2 Error |
|---|---|---|---|---|---|---|---|---|---|---|---|
| **ACE2-Fc** | G614 S | 2.56E-08 | 2.73E-09 | 7.43E+04 | 1.28E+01 | 3.85E+03 | 1.03E+01 | 1.90E-03 | 1.09E-01 | 1.78E-04 | 8.61E-02 |
| **(RBD)** | BA.2 S | 5.68E-09 | 2.44E-10 | 8.31E+04 | 1.28E+01 | 1.69E+03 | 2.29E+01 | 4.72E-04 | 6.18E-01 | 1.79E-05 | 1.10E+00 |
| **Monomeric** | G614 S | 1.54E-08 | 9.83E-10 | 3.42E+05 | | 2.11E+04 | | 5.26E-03 | | 8.98E-05 | |
| **ACE2 (RBD)** | BA.2 S | 2.50E-09 | 6.03E-11 | 9.22E+04 | | 4.06E+02 | | 2.31E-04 | | 5.47E-06 | |
| **C63C8** | G614 S | 6.64E-09 | 2.63E-11 | 1.23E+05 | | 3.97E+02 | | 8.15E-04 | | 1.86E-06 | |
| **(RBD-1)** | BA.2 S | 3.57E-08 | 7.77E-09 | | | | | | | | |
| **G32R7** | G614 S | 3.59E-10 | 6.94E-12 | 1.55E+05 | | 3.88E+02 | | 5.56E-05 | | 1.06E-06 | |
| **(RBD-1)** | BA.2 S | 1.52E-09 | 2.22E-11 | 9.31E+04 | | 5.57E+02 | | 1.42E-04 | | 1.88E-06 | |
| **G32B6** | G614 S | 6.34E-10 | 8.13E-12 | 2.46E+05 | 8.47E-01 | 1.32E+03 | 9.01E-02 | 1.56E-04 | 4.39E-02 | 1.82E-06 | 4.36E-03 |
| **(RBD-2)** | BA.2 S | N.D. | N.D. | N.D. | N.D. | N.D. | N.D. | N.D. | N.D. | N.D. | N.D. |
| **C12A2** | G614 S | 1.08E-09 | 1.38E-11 | 2.03E+05 | 7.29E-01 | 1.30E+03 | 1.24E-01 | 2.19E-04 | 5.68E-02 | 2.42E-06 | 9.11E-03 |
| **(RBD-2)** | BA.2 S | N.D. | N.D. | N.D. | N.D. | N.D. | N.D. | N.D. | N.D. | N.D. | N.D. |
| **C63C7** | G614 S | 5.88E-09 | 2.91E-11 | 1.36E+05 | | 5.80E+02 | | 7.99E-04 | | 1.99E-06 | |
| **(RBD-3)** | BA.2 S | 4.41E-09 | 1.72E-11 | 1.56E+05 | | 4.97E+02 | | 6.90E-04 | | 1.55E-06 | |
| **C12C9** | G614 S | 4.35E-09 | 1.16E-10 | 3.36E+04 | | 6.67E+02 | | 1.46E-04 | | 2.62E-06 | |
| **(NTD-1)** | BA.2 S | N.D. | N.D. | N.D. | | N.D. | | N.D. | | N.D. | |
| **C83B6** | G614 S | 5.70E-10 | 9.19E-12 | 1.27E+05 | | 3.02E+02 | | 7.23E-05 | | 1.15E-06 | |
| **(NTD-1)** | BA.2 S | N.D. | N.D. | N.D. | | N.D. | | N.D. | | N.D. | |
| **C81D6** | G614 S | 9.42E-09 | 3.74E-10 | 5.30E+04 | | 1.55E+03 | | 5.00E-04 | | 1.34E-05 | |
| **(NTD-2)** | BA.2 S | 8.09E-09 | 4.19E-09 | | | | | | | | |
| **C163E6** | G614 S | <1E-12 | 4.16E-12 | 4.50E+04 | | 6.85E+02 | | <1.0E-07 | | 1.87E-07 | |
| **(S2-2)** | BA.2 S | 5.56E-09 | 1.75E-10 | 5.00E+04 | | 8.27E+02 | | 2.78E-04 | | 7.45E-06 | |

**Table S2. Neutralization of the SARS-CoV-2 Omicron subvariants**

| Antibody/ACE2 construct | Epitope | Neutralization titer (μg/ml) | | | | | |
|---|---|---|---|---|---|---|---|
| | | G614 (B.1) | | Omicron BA.1 | | Omicron BA.2 | |
| | | $IC_{50}$ | $IC_{80}$ | $IC_{50}$ | $IC_{80}$ | $IC_{50}$ | $IC_{80}$ |
| C63C8 | RBD-1 | 12.578 | 33.721 | >50 | >50 | >50 | >50 |
| G32R7 | RBD-1 | 0.089 | 0.643 | 0.243 | 1.386 | 0.835 | 3.166 |
| C12E7 | RBD-1 | 3.747 | 30.803 | >50 | >50 | 25.880 | >50 |
| G32B1 | RBD-1 | 8.153 | 41.011 | >50 | >50 | 39.301 | >50 |
| C12A2 | RBD-2 | 0.026 | 0.094 | >50 | >50 | >50 | >50 |
| G32B6 | RBD-2 | 0.016 | 0.056 | >50 | >50 | >50 | >50 |
| C98C7 | RBD-2 | 0.005 | 0.019 | 1.049 | 4.778 | 1.566 | 8.287 |
| G32A4 | RBD-2 | 0.005 | 0.022 | >50 | >50 | >50 | >50 |
| C63C7 | RBD-3 | 10.494 | >50 | 48.823 | >50 | >50 | >50 |
| G32Q4 | RBD-3 | 0.101 | 0.632 | 4.890 | 25.784 | 42.853 | >50 |
| C12C9 | NTD-1 | 0.066 | >50 | >50 | >50 | >50 | >50 |
| C83B6 | NTD-1 | 0.288 | >50 | >50 | >50 | >50 | >50 |
| C93D6 | NTD-1 | 0.778 | >50 | >50 | >50 | >50 | >50 |
| C81D6 | NTD-2 | >50 | >50 | >50 | >50 | >50 | >50 |
| C81G9 | NTD-2 | 12.231 | >50 | >50 | >50 | >50 | >50 |
| C7A9 | S2-1 | >50 | >50 | >50 | >50 | >50 | >50 |
| C163E6 | S2-2 | >50 | >50 | >50 | >50 | >50 | >50 |
| ACE2-T27W-Fd | RBD | 0.098 | 0.468 | 0.036 | 0.126 | 0.031 | 0.084 |
| Positive serum pool_2 (1/x) | N/A | 1,172 | 213 | <20 | <20 | <20 | <20 |
| Normal Human Serum (1/x) | N/A | <20 | <20 | <20 | <20 | <20 | <20 |

### Table S3. Cryo-EM statistics.

| **EM data collection and reconstruction statistics** | | | |
|---|---|---|---|
| Protein | Full-length S protein of the BA.2 variant | | |
| Microscope | Titan Krios | | |
| Voltage(kV) | 300 | | |
| Detector | Gatan K3 | | |
| Magnification(nominal) | 105,000 | | |
| Energy filter slit width (eV) | 20 | | |
| Calibrated pixel size (Å/pix) | 0.83 | | |
| Exposure rate ($e^-$/pix/sec) | 13.761 | | |
| Frames per exposure | 50 | | |
| Total electron exposure ($e^-$/Å$^2$) | 53.853 | | |
| Exposure per frame ($e^-$/Å$^2$) | 1.077 | | |
| Defocus range (μm) | -0.5,-2.2 | | |
| Automation software | SerialEM | | |
| # of Micrographs used | 34,833 | | |
| Particles extracted | 6,763,049 | | |
| Particles after 2D classification | 2,749,398 | | |
| Class | Three-RBD-down | One-RBD-up | One-RBD-inter |
| Total # of refined particles | 160,774 | 193,655 | 113,667 |
| Symmetry imposed | C3 | C1 | C1 |
| Map sharpening B-factor | -87.2 | -79.7 | -72.1 |
| Map resolution (Å) | 2.8 | 3.0 | 3.1 |
| FSC threshold(Å) | 0.143 | 0.143 | 0.143 |
| Model refinement and validation statistics | | | |
| PDB | | | |
| Composition | | | |
| Amino acids | 3384 | 3372 | 3372 |
| Glycans | 57 | 57 | 57 |
| RMSD bonds (Å) | 0.014 | 0.014 | 0.014 |
| RMSD angles (◦) | 1.85 | 2.05 | 2.07 |
| Mean B-factors | | | |
| Amino acids | 95 | 95 | 95 |
| Glycans | 139 | 138 | 139 |
| Ramachandran | | | |
| Favored (%) | 92.04 | 90.88 | 90.72 |
| Allowed(%) | 6.89 | 7.39 | 7.37 |
| Outliers(%) | 1.07 | 1.73 | 1.91 |
| Rotamer outliers (%) | 4.55 | 10.29 | 9.61 |
| Clash score | 2.51 | 5.32 | 5.25 |
| C-beta outliers (%) | 0.95 | 1.65 | 1.30 |
| CaBLAM outliers (%) | 3.32 | 4.41 | 4.05 |
| CC (mask) | 0.81 | 0.80 | 0.77 |
| CC (volume) | 0.80 | 0.79 | 0.77 |
| MolProbity score | 2.02 | 2.58 | 2.56 |
| EMRinger score | 3.42 | 3.23 | 2.99 |
